# Category-preferring patches in human orbitofrontal cortex

**DOI:** 10.1101/2025.04.22.650059

**Authors:** Jianghao Liu, Laurent Cohen, Christophe Pallier, Stanislas Dehaene, Minye Zhan, Paolo Bartolomeo

## Abstract

The human ventral occipitotemporal cortex (VOTC) contains multiple category-preferring areas, organized along posterior-to-anterior and medial-to-lateral axes. Whether this organization extends beyond the VOTC remains unclear. Using two millimeter-scale 7T fMRI datasets, we identified face-and color-preferring patches in the lateral orbitofrontal cortex (OFC) in most participants, and word-preferring patches in half, replicating across runs. As in the VOTC, OFC face and word patches lay lateral to color patches, and OFC patches were as selective as the most anterior VOTC patches. Perception and mental imagery evoked opposite activation gradients across patches, and distinct connectivity patterns, with OFC-anterior temporal coupling persisting during imagery. In congenital aphantasia, OFC selectivity and representational content were comparable to those of typical imagers, but OFC coupling with anterior temporal cortex was weaker. These results suggest OFC and VOTC category-preferring patches belong to shared, functionally connected networks engaged by both visual input and memory.

**One Sentence Summary:** The human orbitofrontal cortex contains category-preferring patches for faces, colors and written words that mirror the topography of ventral occipitotemporal patches and are engaged during both perception and mental imagery.

## Introduction

The ability of object recognition in humans is supported by the ventral occipitotemporal cortex (VOTC)^1^. Within VOTC, multiple sets of areas were found to be sensitive to specific object categories or visual features, including faces^2^, bodies^3^, words^4^, colors^5^, places^6^. In this article, we use “categories” to denote these objects or features, and “preferring” for regions responding more strongly to one category than to the others, without implying exclusive selectivity.

Spatially, these VOTC category-specific areas are hierarchically organized, forming topographic maps that follow two major organizational rules across the cortical space. In the posterior-to-anterior axis from early visual cortices to the anterior temporal cortex, each set of category-specific areas exhibited increasing specificity and invariance, implementing increasing categorical abstraction^1,7^. In the medial-to-lateral axis, regions specific for places, colors, faces, and letter strings were found laying out in this order, which is thought to follow a larger organizational rule of eccentricity^1,8^, where preference for more foveal stimuli would lie in more lateral regions. Most categories are associated with multiple patches spread along the posterior-to-anterior axis, for example, there are parallel series of face- and color-preferring patches in both humans and monkeys^9,10^, and multiple patches sensitive to faces, bodies, spiky, stubby objects in monkeys^11^.

Most previous studies focused on VOTC regions, particularly around the fusiform gyrus and nearby areas, with the anterior temporal cortex considered the apex of these category-specific processes^12^. Category-specificity beyond the VOTC^13,14^ is much less explored. A few studies reported prefrontal face-specific areas (see Ref.^15^ for a review), but it remains uncertain whether the prefrontal cortex contains patches for other categories, and whether those patches show a systematic topographical organization. The orbitofrontal cortex (OFC), as a prefrontal region with direct anatomical connections to visual and anterior temporal areas^16,17^, is a plausible candidate for such a higher-level extension of the visual categorization system. It has also been implicated in top-down visual prediction^18,19^ and in the integration of perceptual information into more abstractly structured cognitive maps^20,21^. Whether its functions are organized according to visual categories remains unclear.

We recently identified category-specific patches for faces and colors in the lateral OFC using high-resolution (1.2 mm isotropic) 7T fMRI during perception and mental imagery of the same items, across five categories (Dataset 1; faces, colors, words, shapes, and maps)^22^. These patches were observed during both perception and imagery in the majority of participants, including individuals with aphantasia who reported a lifelong absence of voluntary imagery^23^. Here, we used this dataset together with an independent 7T fMRI dataset of word reading (Dataset 2^7^) to systematically characterize category specificity in the OFC. In Dataset 1, we examined the topography, category specificity, activation amplitude, representational content, and functional connectivity of OFC face-and color-preferring patches. The combination of perception and mental imagery allowed us to examine OFC responses with and without direct visual input^24–27^, as well as differences between typical imagers and individuals with aphantasia. In Dataset 2, we tested whether OFC category sensitivity extends to written stimuli, identifying OFC word-preferring patches, defined by their sensitivity to the word-similarity effect, in 10 of 21 participants. Across the two datasets, we show that OFC color patches lie medial to face and word patches, paralleling the organization of the VOTC. These findings suggest that category-preferring patches in the OFC and VOTC belong to shared, distributed networks, which can be engaged both by visual input and by content retrieved from memory during mental imagery.

## Results

We report 7-Tesla fMRI individual-subject data, focusing on OFC face- and color-preferring patches (Dataset 1) and OFC word-preferring patches (Dataset 2). Each individual participant serves as a replication unit, by defining patches on one run and testing them on independent runs. We further tested the population prevalence. No spatial smoothing was applied to any individual-level data in this study. We describe the functional anatomy and activation profiles of the OFC patches, as well as their relationship with the VOTC patches. In Dataset 1, we also examine their functional properties, and compare imagery with perception, and aphantasia with typical imagery.

### Dataset 1: Adjacent face- and color-preferring patches in the OFC

Twenty participants (10 typical imagers and 10 aphantasic individuals) compared pairs of stimuli from five different categories, during both visual perception and visual mental imagery: object shapes, object colors, written words, famous faces, and locations of cities on a map of France^22^. The imagery task was carried out before the perception task. For the face imagery task, participants heard a word indicating a category (e.g., “face”), followed by two names (e.g., “E. Macron”, “F. Hollande”). They attempted to conjure and maintain mental images of each face (two names presented 4 s apart; the attribute word was presented 8 s after the second name), then indicate which face best matched an attribute word (e.g., “round”) by pressing a button. For the color imagery task, they decided which of two food items had a darker or lighter color. In the corresponding perception tasks, the same stimuli were presented simultaneously as auditory words and visual pictures. A comparable task using only abstract words, hence minimizing mental imagery, was included as a control. Behavioral results showed that aphantasics performed the imagery tasks as accurately as typical controls, but with slower response times^22^, consistent with the results from larger samples performing the same tasks^28^.

Along the VOTC, we identified individual patches activated by the perception of faces or colors relative to stimuli from the other categories (functional voxels of 1.2 mm isotropic, no smoothing; Region-Of-Interest (ROI) selection: voxelwise p<0.001 uncorrected, cluster size threshold = 12 voxels). For face-preferring patches, which are generally bilateral, the corresponding left- and right-hemispheric patches were merged in subsequent ROI analyses. We identified the following areas: occipital face area (OFA; group-average Talairach coordinates of individual means, with absolute x values: x = 38, y = -79, z = -6), fusiform face area 1 (FFA1; x = 37, y = -63, z = -11), fusiform face area 2 (FFA2; x = 37, y = -47, z = -15), anterior face patch (AFP; x = 36, y = -23, z = -16). We similarly identified bilateral color-preferring patches: posterior color patch (PC; x = 29, y = -69, z = -10), central color patch (CC; x = 30, y = -51, z = -12), anterior color patch (AC; x = 29, y = -34, z = -15). The medial-lateral layout between VOTC color and face patches (Bayesian paired t-test of the absolute x-coordinates, Bayes factor [BF] = 265.76) was consistent with previous work^9,29^. Critically, in the OFC, we also found adjacent face- and color-preferring patches bilaterally during perception. We identified OFC face patches (OFC-f, x = 33, y = 30, z = -6) in 18 out of 20 individuals (all typical imagers and 8 aphantasics) and OFC color patches (OFC-c, x = 26, y = 33, z = -8) in 19 out of 20 individuals (9 typical imagers and 10 aphantasics), showing strong between-subjects replicability. To test replicability of prevalence, patches were defined within-subject in one run (OFC-f: 16/20 in both runs; OFC-c: 13 or 14/20 in run4 and run5, respectively) and tested on the other *independent* run. Both face and color patches showed significant category preference in the held-out run in the majority of participants, with estimated population prevalence (maximum a posteriori) of 0.79 for OFC-f and 0.63 or 0.68 for OFC-c (see Fig. S2, Table S1). Having established their replicability, we retained the OFC patches defined from both perception runs for subsequent analyses, to maximize sensitivity. We did not repeat the cross-validation for VOTC patches, which are well established and showed much stronger responses than OFC. As ROIs were defined from perception trials, all imagery analyses were performed on data independent of ROI definition. Subsequent analyses focused on comparing the functional properties of OFC patches with those of VOTC patches.

The locations of these OFC patches corresponded to specific macroanatomical landmarks, including the caudal parts of both the lateral orbital sulcus (LOS) and the medial orbital sulcus (MOS), and the transverse orbital sulcus (TOS) (Fig. 1A, see inset for schematics of the OFC anatomy, and location occurrences of patches across participants). Color patches were located more medially than face patches in the OFC (Bayesian paired t-test of absolute x-coordinates, BF = 12.68), similar to their medial-lateral layout in the VOTC. The y and z coordinates did not differ between OFC face and color patches in either typical or aphantasic individuals (Bayesian paired t-tests, BFs < 0.92).

**Figure 1.**
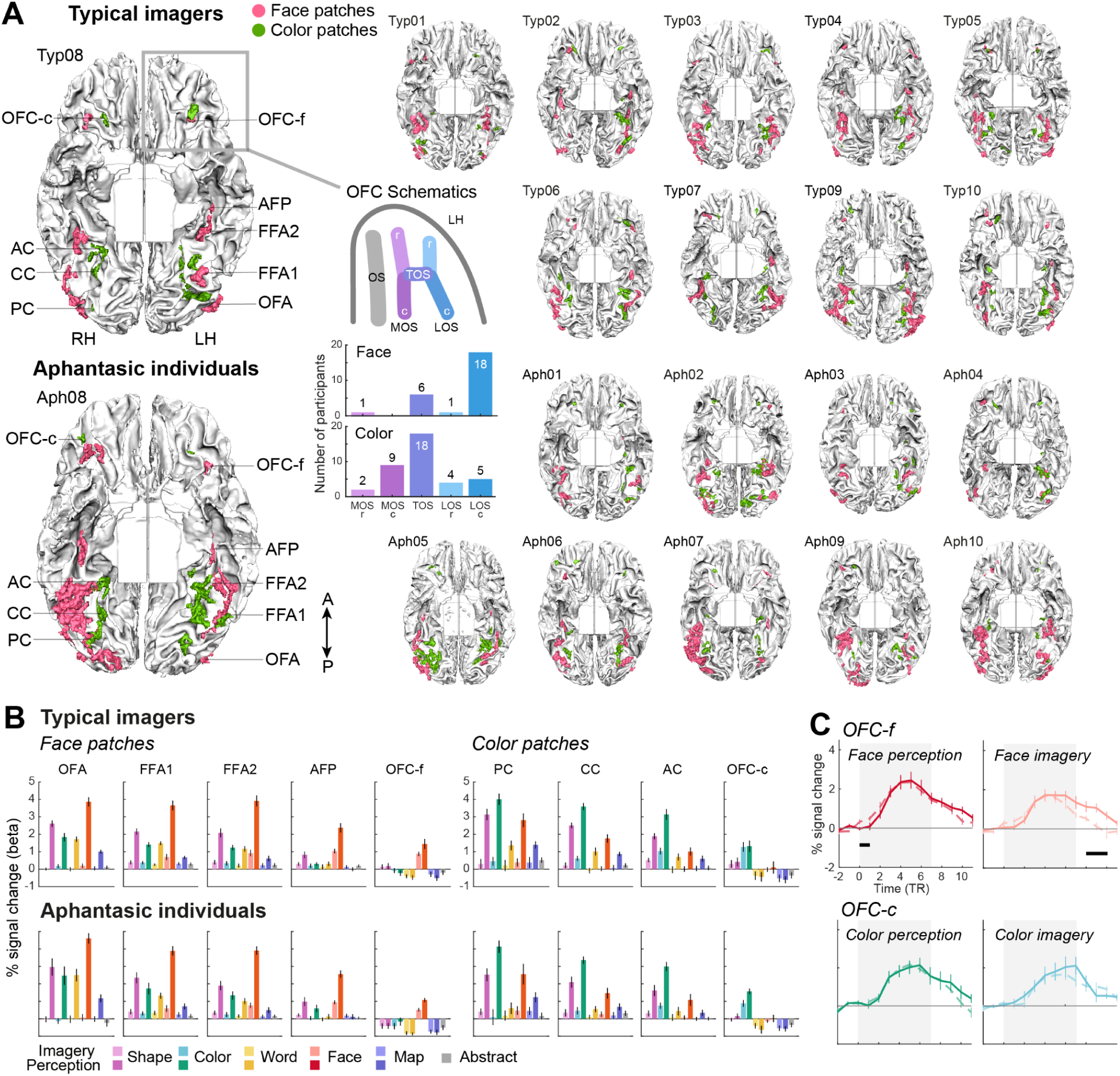
Locations, activity profiles, and time courses of OFC face- and color-preferring patches. **A.** Face and color patches in OFC and VOTC for each individual participant (n=20, including 10 typical imagers [Typ] and 10 aphantasic participants [Aph]). Patches were defined from perceptual trials in the volume space (face or color versus the remaining four categories)^22^, 7T fMRI, 1.2 mm functional resolution, no smoothing, p<0.001 uncorrected), and then rendered onto the segmented white matter surface mesh of each individual participant (ventral view; see Fig. S1 for the rationale of this visualization). From posterior (P) to anterior (A), face patches include: occipital face area (OFA), fusiform face area 1 (FFA1), fusiform face area 2 (FFA2), anterior face patch (AFP), orbitofrontal face patch (OFC-f); color patches include: posterior color (PC), central color (CC), anterior color (AC), orbitofrontal color patch (OFC-c). In perceptual trials, OFC face and color patches were identified in 18/20 and 19/20 individuals, respectively (Aph 01 and Aph 05 lacked face patches, Typ 04 lacked color patches). In most participants, face patches were more lateral than color patches, in both the VOTC and OFC. RH: right hemisphere; LH: left hemisphere. The inset panels show schematics of OFC anatomy^30^ and number of occurrences per OFC sub-region. Most face patches were located in the caudal lateral orbital sulcus (LOS) while most color patches were in the transverse orbital sulcus (TOS) or in the caudal medial orbital sulcus (MOS). No patch was identified in the olfactory sulcus (OS). r: rostral; c: caudal. **B.** Group-averaged activation profiles (percent signal changes) of OFC face and color patches, across five imagery tasks (lighter colors), five perceptual tasks (darker colors) and abstract words task (gray). Error bars represent SEM across participants. Note that OFC patches showed above-baseline activation for preferred categories but b*elow-baseline* activity for most of the other categories. Abstract: a control task with similar structure, but with exclusively abstract words to compare semantic meaning of words (e.g., “routine”, “convention” compared to “society”), to minimize voluntary visual imagery. **C.** Averaged time courses in OFC patches (TR=2 s), plotted in solid lines for typical imagers and dashed lines for aphantasic individuals. Shaded area indicates the period of imagery generation and maintenance, or perception, preceding the response button press. Error bars denote the standard error of the mean across participants. Black horizontal bars represent significant group differences (p < 0.05, Bonferroni-corrected across time points). In OFC face patches, aphantasic participants exhibited higher early perceptual activity and reduced activity during imagery maintenance.

Next, we investigated the general activity profiles of these OFC patches across task conditions using Bayesian ANOVAs with the factors Patch and Task (11 tasks; Fig. 1B), separately for face and color patches. In both typical and aphantasic participants, the BOLD signal changes (beta values) differed between patches and across tasks (BFs for main Patch effects are close to infinity), indicating that activity is not uniform across patches. During perception, the VOTC patches maintained above-baseline activation for non-preferred stimuli, while showing strong activity for their preferred stimuli. However, the OFC patches not only exhibited category-specific activation for faces or colors but also showed below-baseline activity for most of the non-preferred stimuli (Fig. 1B; Bayesian one-sample t-tests, all BFs > 9.49 for visual words, French maps, and abstract words). These patterns were consistent across both groups (see Fig. S3 for activity profiles in individual participants).

In event-related averages across repetition times (TR), OFC activity rose above baseline as early as the first TR after stimulus onset during both perception and imagery (TR=2 s, one-sample t-tests against baseline, all ps < 0.05). The time courses of activity differed between groups in the OFC-f patches (Fig. 1C), showing higher activity in aphantasics during the early phase of face perception (TRs 0 and 1), but lower activity during the maintenance phase of face imagery (TRs 8 to 10) compared to typical imagers. No group difference was observed in OFC-c patches or VOTC patches. Plots for the VOTC patches and all statistics are reported in Fig. S4.

### Category selectivity: posterior-to-anterior increasing trends during both perception and imagery

We quantified the *category selectivity* for each condition relative to other conditions, a critical functional feature of perceptual category-specific areas^2,4^. For each patch, we computed a selectivity index (SI), contrasting activity for the preferred category with the average activity for the remaining four categories: SI = (4 × preferred category − other categories) / (4 × preferred category + other categories), where “other categories” denotes the summed activity of the four non-preferred categories. In electrophysiology studies of faces, activity for faces twice as high as other conditions was deemed “face-selective”^29^, corresponding to a SI of 0.33. To avoid overestimating selectivity due to negative activity amplitudes, negative activity was padded to zero before computing the SI^31^.

Across patches, we observed increases in face- and color-selectivity indices from posterior to anterior patches, during both perception and imagery, and for both typical and aphantasic participants (linear contrast analysis, all ps < 0.003; Fig. 2A). In typical imagers, the SI increased with a slope of 0.13 per patch for face perception (t = 8.37), 0.18 for face imagery (t = 5.92), 0.12 for color perception (t = 4.28) and 0.19 for color imagery (t = 3.31); in aphantasic individuals, by 0.13 for face perception (t = 9.83), 0.21 for face imagery (t = 3.95), 0.15 for color perception (t = 6.76) and 0.33 for color imagery (t = 4.59). There was no evidence for group differences in these trends (Bayesian repeated measures ANOVAs with the factors Group × Patch, interaction effects, all BFs < 0.39; main Group effects, BFs < 0.44, except for color perception, BF = 2.03).

**Figure 2.**
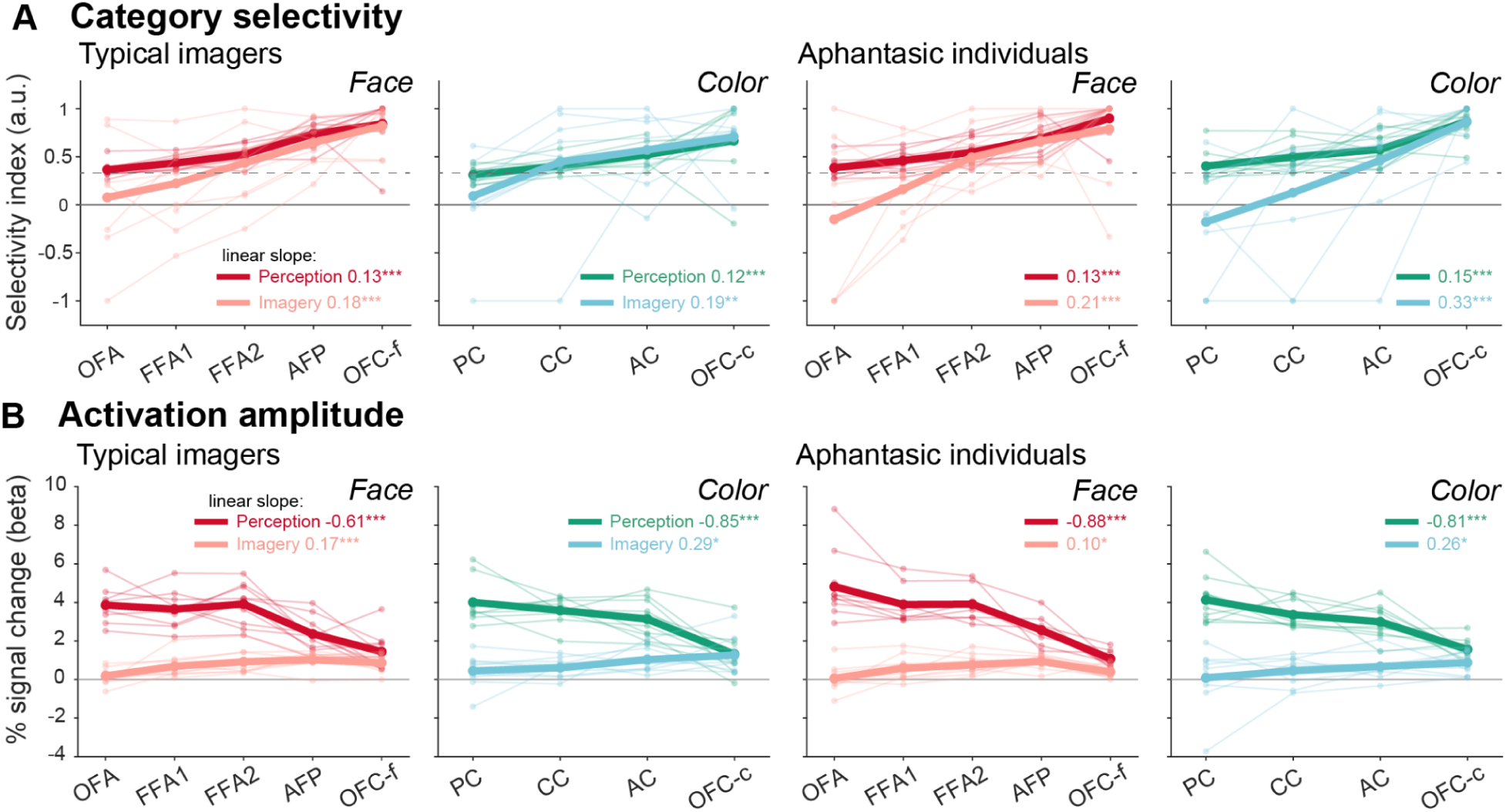
Category selectivity and activation amplitude for face- and color-preferring patches during imagery and perception. In both A and B, bold lines denote the group average, and faint thin lines and dots denote individual participants. Slopes of linear regressions across patches (mean of individual slopes, in units per patch) are shown with the significance of the linear contrast: *p < 0.05, **p < 0.01, ***p < 0.001. The same number of trials of imagery and perception of the same items (in runs 4 and 5) were used for the plot. **A**. Category selectivity increases across VOTC and OFC patches during both perception and imagery. The selectivity index = 0.33 (gray dashed line) corresponds to activity for the preferred category twice that for the other categories. **B**. Across VOTC and OFC patches, the activation amplitude decreases during perception, but increases during imagery.

The OFC patches showed selectivity for their preferred stimuli during both perception and imagery (mean SI = 0.67–0.90) at least as high as that of the anterior temporal patches (AFP and AC, 0.46–0.74), previously considered the most category-selective regions of the ventral visual pathway^1^. OFC selectivity exceeded that of the most anterior VOTC patches during perception (OFC-c vs AC: difference = 0.23, q = 0.020; OFC-f vs AFP: difference = 0.16, q = 0.049), but not reliably during imagery (OFC-c vs AC: q = 0.066; OFC-f vs AFP: q = 0.191; groups pooled, N = 20; two-sided paired t-tests, Benjamini–Hochberg FDR-corrected across the four comparisons).

Because perception estimates are based on the same data used for ROI definition, imagery results provide independent estimates. These OFC patches showed negative selectivity for all non-preferred stimuli (Fig. S5A), confirming their category specificity. Further, cross-validated estimates, in which ROIs were defined on one run and the SI and amplitude computed from the other, confirmed the high selectivity of OFC patches (OFC-f, SI = 0.73; OFC-c, SI = 0.53; both above 0.33, p < 0.01) and their above-baseline responses to the preferred category during both perception and imagery (all ps < 0.01; Fig. S5B, C). Thus their responses were tightly dependent on the stimulus type, but did not depend on the reception of direct visual inputs.

In contrast, the most posterior VOTC patches (OFA and PC) were selective for their preferred category during perception (mean SI = 0.31–0.40), but not during imagery (mean SI = −0.18 to 0.09), meaning that they were not reliably activated by their preferred stimuli in the absence of visual input. Selectivity in these posterior patches did not differ between groups (Bayesian independent samples t-tests, all BFs ≤ 1.02).

### Activation amplitude: opposite posterior-to-anterior trends during perception and imagery

Across patches, we then separately examined the activation amplitudes during perception and imagery. Cross-validated estimates confirmed that OFC patches responded above baseline to their preferred category during both perception and imagery (Fig. S5C), with no group differences. Moving from posterior to anterior patches, including the OFC, the activation amplitudes formed two significant but opposite linear trends for perception and imagery (Fig. 2B). During perception, the activation amplitude decreased from posterior to anterior (linear contrast analysis, slope for face patches = −0.61 % signal change per patch, t = −7.09, p < 0.001; slope for color patches = −0.85, t = −7.94, p < 0.001). Conversely, during imagery the activation amplitude increased monotonically from posterior to anterior (slope for face patches = 0.17, t = 4.68, p < 0.001; slope for color patches = 0.29, t = 2.75, p = 0.011). The same pattern prevailed in aphantasic individuals, showing decreasing slopes in face perception (slope = −0.88, t = −10.76, p < 0.001) and color perception (slope = −0.81, t = −6.36, p < 0.001), and increasing slopes in face imagery (slope = 0.10, t = 2.37, p = 0.023) and color imagery (slope = 0.26, t = 2.34, p = 0.027). The two groups showed comparable activation amplitude (Bayesian repeated measures ANOVAs with the factors Group × Patch, main Group effects, all BFs < 0.62; interaction effects, all BFs < 0.56).

Thus, from posterior VOTC to OFC patches, category selectivity and activation amplitude changed monotonically along the posterior-to-anterior axis, forming functional gradients that extended from the VOTC into the OFC, with opposite amplitude gradients for perception and imagery.

### Multivariate representational content during perception and imagery

Previous studies have shown that some visual areas in aphantasia are activated during imagery, but that their representations may not take a perceptual-like format^22,32,33^. While coding format is hard to assess from mesoscopic fMRI measurements, here we use representational similarity analysis (RSA^34^) to assess whether neural representations retrieved from memory during imagery resemble the feedforward representations evoked by perception, and whether the representational content in OFC and VOTC patches is more closely related to abstract category information or to visual features.

We assessed the overlap between imagery and perception representations by correlating their neural representational dissimilarity matrices (RDMs) for the same items within each patch (Bayesian one-sample t-tests against zero), and comparing the RDMs between groups. In typical imagers, we observed representation overlap in some anterior VOTC patches (Fig. 3A; FFA2 for faces, BF = 4.96; AC for object colors, BF = 9.21). In contrast, no overlap was observed in aphantasia in any patches.

**Figure 3.**
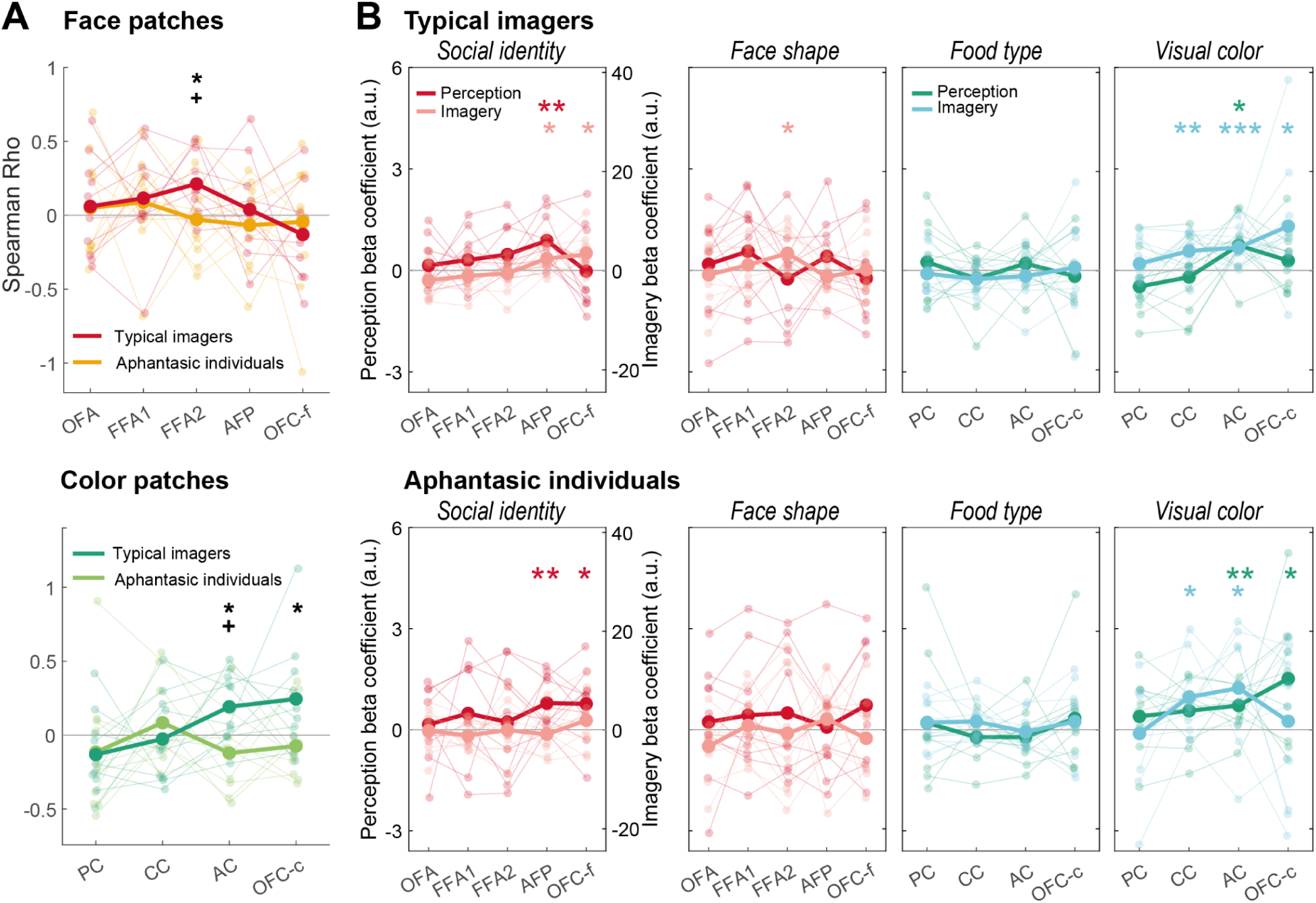
Neural representations. **A.** Representational overlap, quantified as the correlation between imagery and perception neural RDMs. The neural RDMs were computed with the same items from perception and imagery trials in the run 4 and 5. * denotes group difference (BF > 3). “+” denotes above-zero overlaps for BF > 3 in typical imagers. In typical imagers, we observed representational overlap between imagery and perception in some of the anterior regions. No significant overlap was observed in aphantasic individuals in any of these patches. **B.** Representational content. In each patch, the neural RDM was modelled as a linear combination of two behavioral RDMs, resulting from the participants arrangement of fMRI stimuli based on either their semantic or visual features in a behavioral session (Fig. S6). Plots represent the beta coefficients from those models. Face RDMs correspond to either the celebrities’ social identity (profession), or their face shapes; color RDMs correspond to food types (semantic content of food stimuli), or the visual color appearance. Bold lines represent group averages, and faint lines represent individual participants. Red/pink lines denote face trials, and green/blue lines denote color trials. *BF > 3, **BF > 10, ***BF > 30, Bayesian one-sample t-test versus 0, indicating the presence of representational content. The scales of beta coefficients differ between perception and imagery, as the perceptual results were derived from 6-item RDMs, and the imagery results were derived from 18-item RDMs.

Compared to typical imagers (Bayesian independent samples t-test), aphantasic individuals showed a reduction in overlap in OFC-c (BF = 3.87), FFA2 (BF = 3.08), and AC (BF = 7.71) patches.

In a post-fMRI behavioral session, we assessed participants’ individual mental representations of the fMRI stimuli, and correlated those patterns to the neural representations within cortical patches. Participants were instructed to arrange the same stimuli presented in the fMRI experiment inside a circular region, based either on their semantic or their visual features. For faces, participants organized the written names of the celebrities according to the core social marker of the profession, and arranged celebrities’ photos according to the shape of their face (round or elongated). For colors, participants categorized written names of food items according to their semantic proximity (vegetables, tropical fruits and processed food), and arranged food photos according to their color. We derived behavioral RDMs based on individual participants’ arrangements (Fig. S6 displays the arrangements per condition, for the 18 items from the fMRI imagery trials). Neural RDMs were derived by calculating pairwise Pearson correlations between multi-voxel patterns of stimuli during perception or imagery. For each experimental condition, we predicted the neural RDM of each patch using the semantic and visual behavioral RDMs of the corresponding items as linear predictors.

The behavioral RDMs revealed no significant differences between typical and aphantasic participants (all ps > 0.09), indicating comparable representational content across groups. Similarly, there were no group differences in the neural representations observed in any patch or condition (Bayesian independent samples t-test, all BFs < 2.65).

Stimulus-specific content was represented in the OFC and anterior VOTC patches but not in the posterior patches (Fig. 3B, Bayesian one-sample t-tests against zero in each patch). Most of these patches were sensitive to the social identity of faces, or to the visual colors, although the conditions in which these effects reached significance varied across patches and groups. Face shape information was only present in FFA2 for typical imagers during imagery (BF = 4.51). No evidence was found for food-related semantic representation in the color patches of either group (all BFs < 0.63).

Among OFC patches, the OFC-f coded social features (in typical imagers, during imagery, BF = 5.77; in aphantasics, during perception, BF = 3.38). The OFC-c patches coded visual color (in typical imagers, during imagery, BF = 3.26; in aphantasics, during perception, BF = 4.98). The OFC patches thus carry behaviorally relevant neural information during both imagery and perception, with moderate evidence.

Among VOTC patches, the AFP coded social identity during both face perception (typical imagers, BF = 20.06; aphantasics, BF = 11.23) and face imagery (typical imagers, BF = 5.78). The AC coded visual color during both color perception (typical imagers, BF = 7.56; aphantasics, BF = 12.33), and color imagery (typical imagers BF = 36.98, aphantasics BF = 4.01). The visual color representation was additionally found in the CC during imagery for both groups (typical imagers, BF = 10.14; aphantasics, BF = 7.98).

In summary, the OFC patches—like the anterior temporal patches—encode stimulus-specific content, with no evidence for differences between groups. However, only typical imagers exhibited perceptual-like representation during imagery.

### Functional connectivity: opposite patterns during perception and imagery

We examined task-modulated functional connectivity by psychophysiological interaction (PPI) analysis. This method explores the co-fluctuated activity in the OFC patches and VOTC patches, as well as the modulation between them during both perception and imagery tasks. This allowed us to determine whether these patches belong to a connected functional network. Although the method assesses only the co-fluctuation of fMRI signals, we adopted the term “functional connectivity” which is by now classical in the literature.

We computed the pairwise connectivity between the OFC, VOTC patches, and additionally to V1, contrasting the perception and imagery tasks separately relative to the abstract-words control task (comparison of beta values versus 0 with Bayesian one-sample t-tests for the pairwise connectivity between regions), and compared the resulting maps between perception and imagery. In typical imagers, the same pattern prevailed for faces and colors. The functional connectivity between the OFC-f and AFP patches was higher during face imagery compared to face perception (Bayesian pairwise t-test, BF = 5.87). There were distinct connectivity patterns during imagery and perception. During perception, we observed strong and wide-spread functional connectivity between the V1, VOTC, and OFC patches (compared to the control task, see detailed pairwise connectivity strengths in Fig. 4A). Consistent with this ROI-based result, a complementary whole-brain seed-based analysis showed that, during perception, OFC-f and OFC-c were functionally coupled with anterior VOTC regions near the corresponding category-specific patches, and OFC-f additionally with the amygdala (Fig. S7).

**Figure 4.**
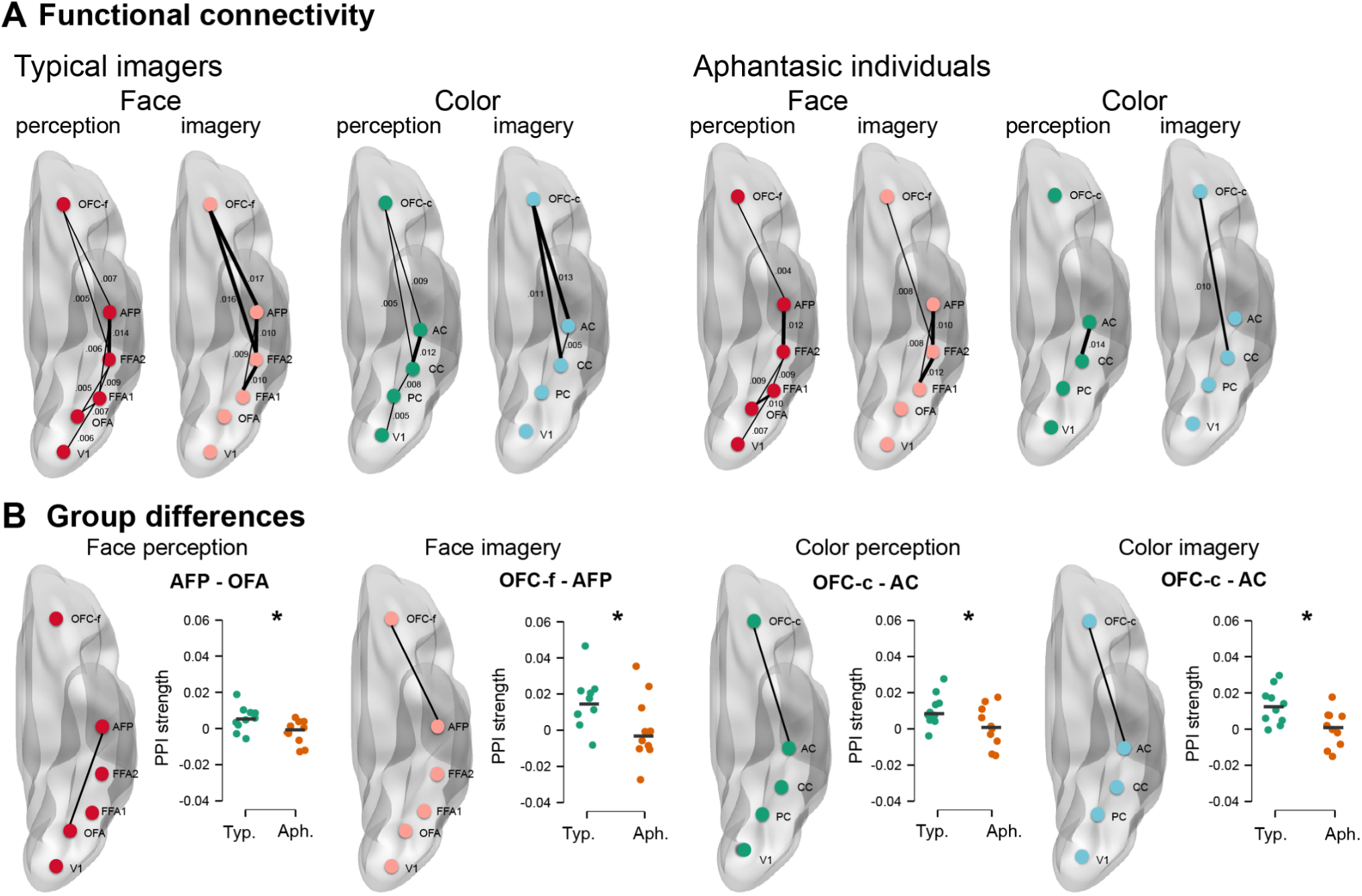
Task-dependent functional connectivity during imagery and perception. **A.** In typical imagers, the strength of PPI modulation, contrasting perception or imagery to an abstract words control condition. Lines indicate significant connectivity between two patches, as tested by Bayesian one-sample t-test against zero. Line thickness and values indicate group-level PPI strength. **B.** Group differences in PPI strength. Lines indicate higher connectivity in typical imagers than in aphantasic individuals. Plots show connectivity strength in individual participants. Horizontal bars indicate the group average. * denote significant group differences (BF > 3). Typ, typical imagers; Aph, aphantasic individuals.

In contrast, during imagery, V1 and the posterior VOTC patches were not significantly connected, while the OFC and the anterior VOTC patches remained strongly interconnected. In aphantasic individuals, we observed across-area connectivity patterns similar to those of typical imagers during perception, but aphantasics appeared to have fewer pairs of functionally connected patches during imagery (Fig. 4A).

Comparing the functional connectivity between groups revealed lower connectivity in aphantasics than in typical imagers, mostly involving the OFC (Bayesian independent samples t-test, separately during perception and imagery). Aphantasic individuals had lower connectivities between the anterior patches OFC-f and AFP during face imagery (Fig 4B, BF = 3.27), and between OFA and AFP during face perception (BF = 5.39). For color patches, aphantasics had lower connectivity between the OFC-c and AC patches during both color perception (BF = 3.00) and imagery (BF = 8.18).

### Dataset 2: Word-preferring patches in the OFC

The color and face patches in the OFC followed a medial-lateral layout similar to that in the VOTC. This suggests that a topographic map of multiple category-specific patches may be a shared feature of both the OFC and VOTC. Besides face and color patches, patches for other categories (e.g. words, bodies, places) might also be present in the OFC. In Dataset 1, however, category-preferring patches for words, shapes and maps were mostly found in the posterior-to-middle fusiform gyrus or in the collateral sulcus; no reliable patches emerged in the anterior temporal cortex or OFC. This may reflect the limited sensitivity of Dataset 1 for these categories, with only six perceptual trials per category.

We therefore assessed the presence of OFC word patches using an independent 7T fMRI dataset^7^, with a design optimized for word processing, in which 21 participants passively viewed 14 conditions of six-letter strings that varied in the frequency of word components (letters, bigrams, and quadrigrams) in English or French, with 15 repetitions per condition. This dataset previously showed that VOTC word-preferring patches, independently defined using a passive-viewing localizer (Fig. 5A, blue patches), exhibited a word-similarity effect in the main experiment, whereby letter strings that were more similar to real words elicited greater activity, as characterized by a significant linear-regression slope across the 14 conditions. This word-similarity effect, which increased from posterior to anterior VOTC, provides a sensitive functional signature of word-selective regions, particularly in more anterior regions where conventional word-selectivity contrasts may be less sensitive. We therefore used this effect to directly assess the presence of word-preferring patches in the OFC (Fig. 5A, yellow patches).

**Figure 5.**
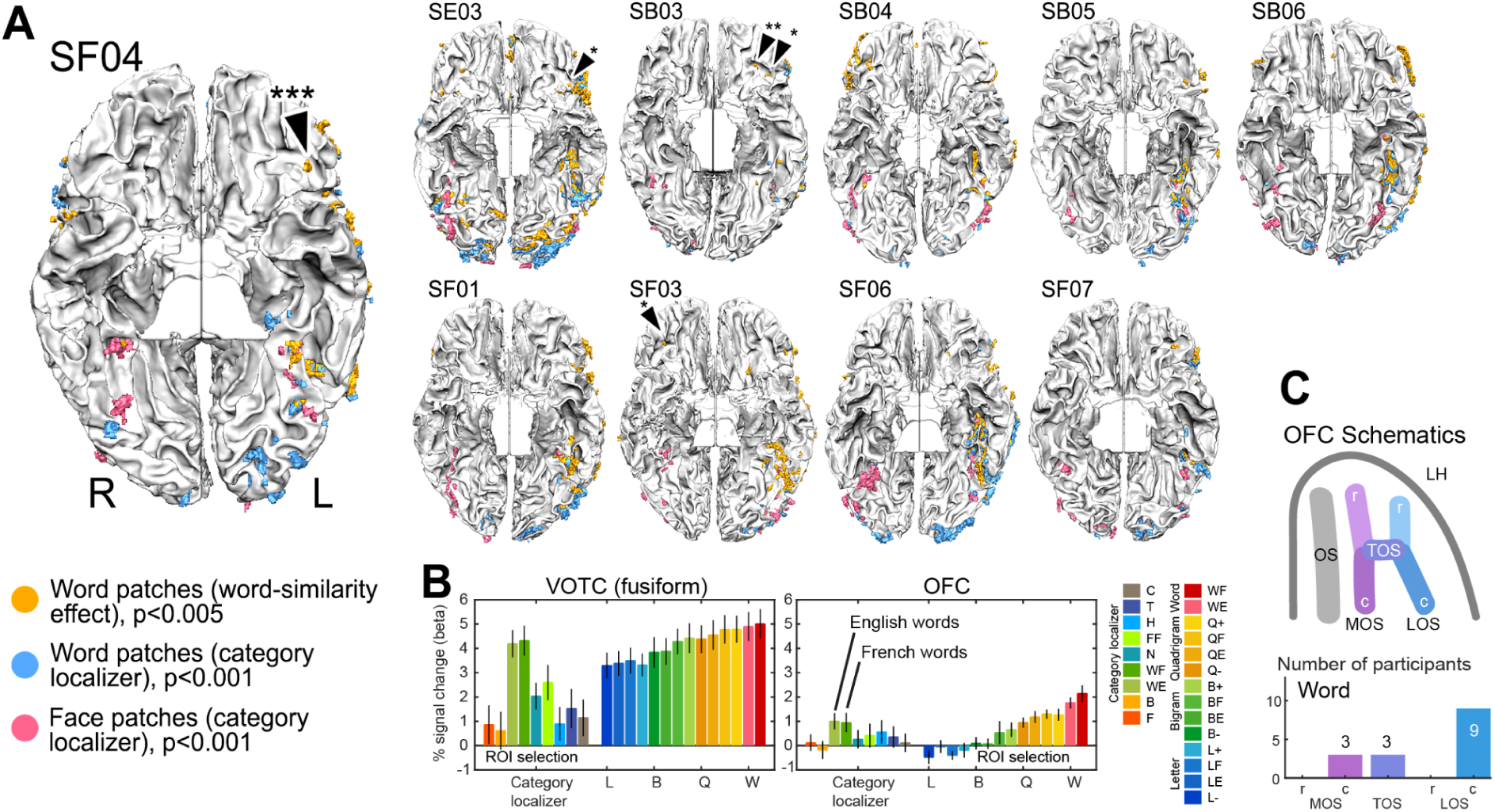
Individual word-preferring patches in the OFC and VOTC. **A.** Word-preferring patches in 10 out of 21 participants (bilateral in 2 participants, left-hemisphere-only in 7 participants, right-hemisphere-only in 1 participant.). The OFC patches were observable when defined by the word-similarity effect (yellow patches, 15 repetitions per condition), and were far less observable in an independent object category localizer (blue patches, 5 repetitions per condition, Table S2). Face-preferring patches (pink) defined by the localizer were plotted to show the medial-lateral organization in VOTC. All maps were cluster-size thresholded at > 4 functional voxels. The patches are rendered onto the segmented white matter surface mesh of each individual participant (ventral view, see Fig. S1 for the rationale of this visualization). Black arrowheads indicate the OFC patches replicated by ROI GLM in the localizer data (English and French words > faces, bodies, houses, tools). *p<0.05; **p<0.005,***p<0.001. SE, SB, SF: participants who are English-dominant, English-French early bilinguals, and French-dominant. **B.** For descriptive purposes, the averaged activity profiles across the localizer (left side, 9 conditions) and the main experiment (right side, 14 conditions), in VOTC and in OFC ROIs. Error bars denote SEM across 10 participants. Note that the VOTC ROIs were defined by the localizer data, and the OFC ROIs were defined by the main experiment data, while the remaining conditions per panel were independent from ROI definitions. See Figure S8 for data of individual participants. Abbreviations for conditions in the localizer: F: faces, B: bodies, WE: English words, WF: French words, N: numbers, FF: false fonts, H: houses, T: tools, C: checkerboards; in the main experiment: L: letters, B: bigrams, Q: quadrigrams, W: words, E or F: word component frequency high in only English or only French, - or +: word component frequency low or high in both English and French. **C.** The number of occurrences of OFC word patches across 10 participants.

We performed Spearman correlation between the word-similarity predictor and the beta values per voxel across 14 conditions, and thresholded whole-brain Spearman Rho maps at p<0.005. The word-similarity effect (yellow patches) was not significant in early visual cortices in most participants, consistent with our previous observation^7^.

Within the fusiform regions, patches with a word-similarity effect overlapped with the ones defined by the localizer (Fig. 5A, yellow versus blue patches), and showed a strong left-hemisphere bias (paired t test, left vs right hemispheres: 6.5 vs 1 patches, p=9.93x10^-5^; 248.08 vs 16.1 functional voxels, p=2.4x10^-4^). In some participants, it revealed more anterior patches extending into the anterior ventral temporal cortex.

OFC word patches with a positive word-similarity slope were observed in 10 out of 21 participants (Fig. 5A, p<0.005). As in VOTC regions, OFC activation amplitude tended to increase as letter strings became more similar to real words (Fig. 5B, individual data in Fig. S8; Table S2 and Fig. S9 with different thresholds). One participant showed an OFC patch with negative slope (higher activity to less word-like stimuli, Fig. S8C) and was not included in subsequent analysis apart from cross-run prevalence validation (performed on all participants). In cross-run prevalence validation, OFC word patches also showed medium, significant replicability: when ROIs were defined in *all* 21 participants, and tested by leaving one run out, they showed a significant word-similarity effect in 8-12 of 20 or 21 participants per fold, with an estimated population prevalence of 0.37-0.55 (96% HPDI lower bounds 0.16-0.32, and upper bounds 0.60-0.76; Table S3), and a group-mean ρ significantly above zero in every fold (ρ=0.34-0.40, all ts > 3.90, all ps <0.001; Figure S10).

Similar to the OFC face and color patches, most of the OFC word patches (Fig. 5C) were located in the caudal portion of the LOS (9/10 participants), and much less in MOS (3/10 participants) and TOS (3/10 participants). The number of OFC word patches showed a left-hemisphere bias (paired-sample t tests, 1 versus 0.3 patches, p=0.025), comparable to the VOTC word patches (χ^2^(1)=0.835, p=0.36). The number of voxels was not significantly left-biased (7.46 versus 2.78 voxels, p=0.118). The OFC word patches were located lateral to the OFC color patches of Dataset 1 (group-average absolute Talairach x coordinates, 35 versus 26; Wilcoxon rank sum test for independent samples, p=0.003), but did not differ from the OFC face patches (35 versus 33, p=0.51), although the VOTC word patches specifically around the fusiform gyrus were located lateral to the face patches (Dataset 2 localizer, 40 versus 38, paired-sample t test, p=0.016).

The Dataset 2 localizer had limited sensitivity (passive viewing, five repetitions per condition), with OFC face and word activations observed in very few participants even at more lenient thresholds (Table S2). Nevertheless, in this independent localizer, the OFC word patches defined by the word-similarity effect showed a word SI significantly greater than zero (mean SI = 0.32, one-sample t-test, t(9) = 3.34, p = 0.005), and comparable to that of the fusiform word patches (0.43; paired t-test, p = 0.197), supporting their word preference relative to other categories. Thus, the absence of OFC face patches at the standard threshold in Dataset 2 likely reflects this limited sensitivity, which also revealed the AFP in only a few participants (Fig. 5A), rather than a lack of replicability, as the OFC face patches of Dataset 1 replicated across independent runs (Fig. S2).

## Discussion

Faces, colors, and written words are important classes of visual objects processed by dedicated category-specific regions in the VOTC^2,4,5^. Using two millimeter-scale 7T-fMRI datasets, we show that this functional specialization extends to the lateral OFC, where we identified patches with preference for faces, colors and words. OFC color patches were located more medially to OFC face and word patches, following a layout similar to that of the VOTC, and all OFC patches showed medium to high inter-individual replicability. In Dataset 1, during both perception and imagery, face- and color-preferring patches showed a monotonic increase in category selectivity along the posterior-to-anterior axis, with OFC patches comparable to the most anterior VOTC patches. However, perception and imagery exhibited opposite trends in activation amplitude and distinct patterns of functional connectivity. Representational similarity analysis further provided moderate evidence that the OFC patches encoded behaviorally relevant content, with face patches encoding social identities (professions) and color patches encoding visual colors. In individuals with aphantasia, OFC patches retained their category specificity but showed weaker connectivity with anterior temporal patches, and imagery representations were less perception-like. In Dataset 2, word patches were sensitive to the word-similarity effect, with increasing activity for letter strings similar to real words. Together, these findings suggest that the OFC contains a topographic organization similar to that of the VOTC, and that category-preferring patches in both regions belong to shared, functionally connected networks engaged by both perception and mental imagery. To our knowledge, this is the first systematic characterization of color and word patches in the OFC and of face patches in the human lateral OFC.

### Face-, color- and word-preferring patches in the OFC

We next consider each category in light of previous macaque and human OFC studies, which have mostly focused on single categories.

Face-preferring patches in OFC have been described in non-human primates^35,36^ and occasionally found in humans^12,37,38^, at locations comparable to ours. In monkeys, OFC face neurons were sensitive to views, emotional expressions, and socially relevant facial features such as age and sex^36,39,40^. Our findings are consistent with these proposed functional roles, suggesting that human OFC face patches may represent social identity, such as abstract categories of professions, which are critical for human social cognition.

Anatomical sites consistent with our OFC color-preferring patches have been associated with achromatic food pictures^35,41^ or food-related feature dimensions^42^, suggesting a food-related representation. This possibility may be relevant because the stimuli in our color task consisted of food images presented in different colors. However, food category selectivity alone could not fully explain our findings. First, our OFC color patches represented the visual colors of the stimuli in our task, with no evidence for category-level food representations. Second, our VOTC color patches correspond to conventional color-biased areas^9,43^ activated by a variety of non-food visual stimuli, and in color-biased regions, color explains unique variance in voxel responses beyond food selectivity, including in color-related voxels extending into the OFC^44^. More generally, color may serve as a functional tag for distinguishing objects and their co-occurring scene attributes, as food objects are often associated with warm, saturated colors, whereas backgrounds tend to be cooler^45^. OFC color patches may encode the behavioral relevance of chromatic signals, potentially including food-related evaluation, such as inferring taste or ripeness from color cues^46^.

The lateral OFC has been associated with multimodal experiential representations linked to lexical-semantic processing, including object and event nouns^47^. Consistent with this, a recent 7T study of word and sentence processing identified an OFC patch that may be involved in processing concrete words^48^. Our OFC word-preferring patches were sensitive to high-level orthographic and visual regularities that increased with similarity to real words, as indexed by the word-similarity effect, suggesting that they may encode abstract aspects of word-like structure such as word frequency beyond individual letter features, through which word-like visual forms could engage higher-order lexical or experiential representations. Their preference for words relative to other categories in an independent localizer was comparable to that of fusiform word patches (mean SI = 0.32 versus 0.43), although it should be confirmed with more sensitive localizers.

Across these three categories, a common OFC function may involve the representation and integration of behaviorally relevant information, consistent with the OFC cognitive map theory^21^, which proposes that the OFC represents abstract task states and integrates information about stimuli and their relationships to guide flexible behavior^20,49,50^. Our findings extend this account by showing that such abstract structure can retain a systematic visual category preference. OFC face- and color-preferring patches were highly selective for their preferred categories, and word patches were sensitive to the word-similarity effect and preferred words over other categories, while the representational content of face and color patches captured behaviorally relevant information such as social identity and visual color. Thus, category-preferring representations may constitute structured components of broader OFC cognitive maps transforming categorical sensory or memory contents into behavioral decisions. Future studies could test whether OFC category-preferring patches exhibit grid-like modulation, a neural signature supporting cognitive-map representations^20^, along behaviorally relevant dimensions, such as social hierarchy in face patches^51^ or semantic dimensions in word patches^52^.

### Shared multi-category topography across the OFC and VOTC

Our findings suggest that the processing of visual categories extends beyond the visual cortex into a functionally connected network spanning the VOTC, the OFC, and possibly the LPFC. The OFC shows anatomical and functional properties that mirror those of the VOTC. In non-human primates, the OFC has direct anatomical connections with aIT^53,54^, including selective projections between face patches^55^, and in humans there is OFC-to-VOTC effective connectivity^17^. In addition, the medial-to-lateral arrangement of color and face/word patches in the OFC parallels the organization of the VOTC. In the VOTC, this arrangement has been related to cytoarchitectonic constraints^56^ and to retinotopic eccentricity biases^1,8^. Similarly, the OFC patches appear to align with distinct cytoarchitectonic subdivisions. Among the cytoarchitectonically defined subdivisions of the lateral OFC^57^, area Fo4 overlaps with our color patches and has been associated with gustatory processing in meta-analytic decoding, whereas areas Fo6 and Fo7 overlap with our face and word patches and have been associated with face and language processing. Eccentricity biases in OFC need to be further validated since evidence for a retinotopic organization in OFC is still lacking. A more plausible shared constraint is long-range anatomical connectivity, with white matter pathways linking corresponding patches across the two regions. This is consistent with the proposal that a region’s category preference depends on its connectivity with a network of regions that also process information about that cognitive domain^58^.

Repeated multi-category topographic maps in OFC and VOTC suggest a fundamental organizational motif in this system. Similar to the tightly clustered and repeated multi-category topographies observed in the monkey visual cortex^11^, the repeated multi-category topography observed in the OFC may represent an additional set of nodes beyond the VOTC. This organizational motif may extend further into the LPFC. Previous work has identified single-category-specific areas in ventral LPFC, for faces in both monkeys^36,59^ and in humans^60,61^, and for scenes in humans^62^. A recent study further provided evidence for face/place/body/tool patches distributed surrounding LPFC multiple-demand regions^63^. In our mental imagery dataset^22^, the inferior frontal gyrus also contained category-specific areas functionally connected with relevant fusiform regions during imagery of object shapes, colors, words and faces.

Together, this evidence suggests that category-preferring processing is supported by repeated multi-category maps distributed across the VOTC, OFC and possibly LPFC, whose shared layout may be constrained by cytoarchitecture and by long-range anatomical connectivity between corresponding patches. How these maps interact, and whether their functional properties differ when information is generated internally rather than received from the external world, can be addressed by comparing perception to imagery and aphantasia to typical imagers, as discussed below.

### Top-down dynamics between the OFC and VOTC

Regarding how these maps interact, an influential account proposes that the OFC acts as a “top-down predictor” that facilitates visual recognition^18,19^: it receives rapidly available visual information driven by low-spatial-frequency signals, and generates predictions that constrain the interpretation of ambiguous visual inputs. Consistent with this proposal, OFC responses in humans have been reported to precede those in middle and posterior fusiform regions by approximately 50 ms and to show functional connectivity with visual regions^18^. In macaques^64^, OFC face patches, with anatomical locations and face selectivity similar to ours, responded with short latencies (∼70 ms) comparable to the most anterior temporal face patch, but later than the ventral LPFC face patches (∼40 ms), suggesting that the aIT, OFC, and LPFC may constitute dissociable stages within a single face-processing system, with the prefrontal stages not simply serial successors of the ventral stream. Fast subcortical routes may also contribute, such as the amygdala, which we found strongly activated during face perception and imagery^22^, and which conveys low-spatial-frequency information with very short latencies^65,66^.

Our findings may extend the top-down predictive account to category-specific processing. Our design did not manipulate expectations or measure the timing of responses, and therefore cannot establish the direction of information flow.

Nevertheless, two observations are difficult to reconcile with purely feedforward visual processing. Along the posterior-to-anterior axis, from occipitotemporal to OFC patches, activation during perception decreased, whereas activation during imagery increased, so that the difference between perception and imagery was smallest in the OFC. Functional connectivity showed a similar dissociation: during imagery, the OFC and anterior VOTC patches remained interconnected, whereas V1 and posterior VOTC patches were connected mainly during perception. These opposite gradients are consistent with the view that the visual cortex acts as an adaptive processor, whose responses are shaped by top-down feedback according to the current task^67^. During perception, responses are dominated by feedforward input and decrease as information progresses anteriorly; during imagery, when feedforward input is absent, responses instead increase toward the OFC, consistent with the OFC lying closer to the source of top-down generative signals. Opposite patterns of connectivity during perception and imagery have also been reported between visual areas and other frontal regions^68,69^.

Such patterns are usually interpreted as a reversal of information flow within a single visual hierarchy, but perception and imagery may also enter a heterarchical network at different nodes^27^, with communication between content-specific regions established flexibly according to task demands. Beyond generating predictions from coarse visual input, the OFC may thus convey category-specific information in the absence of external visual input, to be elaborated into a more detailed visual representation in the VOTC.

### Altered OFC activity and connectivity in aphantasia

Comparing typical imagers and individuals with aphantasia may clarify the contribution of the OFC to visual imagery. Aphantasic individuals report greater difficulties with face^70,71^ and color tasks^72,73^, yet retain access to visual knowledge about these categories ^28^. Here, OFC and VOTC patches in aphantasic individuals showed category selectivity, activation trends and representational content comparable to those of typical imagers, suggesting that aphantasia does not reflect an absence of category-specific representations that can be engaged without visual input. However, imagery representations were less perception-like in the OFC-c and anterior VOTC patches (FFA2 and AC), suggesting that category-specific signals may reach these regions without being reinstated in a perceptual format^22,33^.

The most consistent group differences involved the OFC. The OFC-f patches in aphantasic individuals showed higher early activation during face perception but less sustained activation during face imagery (Fig. 1C), and OFC patches showed weaker functional coupling with the anterior temporal patches of the corresponding systems (Fig. 4B). This pattern is consistent with the broader functional disconnection account of aphantasia^74,25,75^, with reduced microstructural integrity of the uncinate fasciculus linking the OFC and anterior temporal cortex^76^, and with the three-stage model of conscious imagery^25^, in which relatively preserved generation is followed by altered integration and amplification. Altered early iconic memory^77^ and recurrent visual processing in visual pattern completion^78^ in aphantasia suggest that weaker OFC-mediated signals could affect these stages, which remains to be tested directly. Given the modest sample size and the moderate strength of evidence (BFs = 3–8), these group differences, which may also partly reflect differences in task strategy or effort, should be further confirmed in larger samples.

### Why OFC category-preferring patches have been difficult to identify

Only a few previous studies have reported OFC patches selective for visual categories, potentially because the OFC was outside the field of view in many human fMRI studies^9^, OFC responses can be relatively weak and noisy^36^, and its category-specific patches are small and anatomically variable across individuals. Moreover, the OFC is highly susceptible to signal dropout. In our data, the identified patches were located in lateral OFC regions with adequate tSNR, comparable to that of VOTC patches and similar between aphantasic and typical imagers (Fig. S11). Candidate patches likely to reflect signals from large blood vessels, to which gradient-echo EPI at 7T is sensitive, were excluded during ROI definition (see Methods). In addition, the detection of OFC patches may depend on how strongly the task and stimuli engage the corresponding category representations. In Dataset 1, no reliable OFC word patches were identified, possibly because the word task, based on judgments of letter shape (ascenders/descenders), engaged only lower-level orthographic features. In Dataset 2, OFC word patches were revealed by a parametric manipulation of word-similarity with 15 repetitions per condition, whereas the standard passive-viewing category localizer (five repetitions per condition) identified face and word patches mostly in the middle fusiform regions, but not reliably in the anterior ventral temporal cortex or OFC. Reliable patch localization for these three and more categories therefore benefits from high-field, high-resolution fMRI with sufficient OFC coverage, sensitive stimulus manipulations with enough repetitions, and individual-subject analyses.

To conclude, our findings suggest that face- and color-preferring patches in the OFC, together with OFC word-preferring patches sensitive to the word-similarity effect: 1) are observed in most participants and replicate across independent runs, with face patches lateral to color patches, as in the VOTC; 2) carry behaviorally relevant attributes (social identity for faces, visual color for colors); and 3) are engaged during both perception and mental imagery, with preserved selectivity but altered connectivity in congenital aphantasia. Their selectivity is comparable to that of the most anterior VOTC patches, suggesting that they are functionally connected nodes of distributed category-specific networks. The OFC patches may thus link category-preferring visual representations with context-dependent cognition, whether the content arrives from the retina or is retrieved from memory.

## Author contributions

Conceptualization: J.L., M.Z., P.B., L.C., S.D., C.P.; Data curation: J.L.; Formal analysis: J.L., M.Z.; Investigation: J.L.; Methodology: J.L., M.Z.; Funding acquisition: J.L., P.B., S.D.; Resources: S.D.; Project administration: P.B., J.L.; Software: J.L.; Supervision: P.B., S.D., M.Z.; Visualization: J.L., M.Z.; Writing - original draft: J.L.; Writing - review & editing: J.L., M.Z., L.C., P.B., S.D., C.P.

## Materials & Correspondence

Requests for materials and correspondence should be addressed to Jianghao Liu.

## Acknowledgements

The research was supported by INSERM, CEA and specific funding from Dassault Systèmes. The work of J.L. is supported by research funding from Dassault Systèmes. The work of P.B. is supported by the Agence Nationale de la Recherche through ANR-16-CE37-0005 and ANR-10-IAIHU-06, and by the Fondation pour la Recherche sur les AVC through FR-AVC-017. The work of M.Z. is supported by Programme Investissements d’Avenir IHU FOReSIGHT (ANR-18-IAHU-01).

## Methods

### 7T fMRI data acquisition

We re-analyzed two previously collected 7-Tesla fMRI datasets^7,22^. Both datasets were acquired with the same MRI sequences and brain coverage on the same machine at NeuroSpin. High-resolution anatomical images were acquired with an MP2RAGE sequence (resolution=0.65 mm isotropic, TR=5000 ms, TE=2.51 ms, TI1/TI2=900/2750 ms, flip angles=5/3, iPAT=2, bandwidth=250 Hz/Px, echo spacing=7 ms). Functional data were acquired with a 2D gradient-echo EPI sequence (TR = 2000 ms, TE = 28 ms, voxel size = 1.2 mm isotropic, multiband acceleration factor=2; encoding direction: anterior to posterior, iPAT=3, flip angle = 75, partial Fourier=6/8, bandwidth=1488 Hz/Px, echo spacing=0.78 ms, number of slices=70, no gap, reference scan mode: GRE, MB LeakBlock kernel: off, fat suppression enabled). Distortion correction runs with the opposite phase-encoding direction were collected for each task run. The acquisition slab covered the majority of the cortex, avoided the eyeballs, and minimized the ear-canal signal dropout, allowing us to observe very anterior temporal patches around the ear-canal region (see Figure 1 and Figure 5 for example patches). The temporal pole and the superior part of the primary motor cortex (foot regions) were outside the coverage. Both studies^7,22^ were approved by CEA and the local ethics committee in the NeuroSpin Center (CPP 100032 and 100055), and the study was carried out in accordance with the declaration of Helsinki.

### Dataset 1: mental imagery and perception

In this dataset^22^, 20 healthy participants (10 typical imagers, 29.28 ± 8.47 years, 6 female and 10 aphantasic individuals, 28.69 ± 8.27 years, 6 female) performed the enhanced Batterie Imagerie-Perception - eBIP^28^. The battery assessed: i) imagery of object shapes, object colors, faces, letters, and spatial relationships on an imaginary map of France; ii) a non-imagery control task using abstract words, and iii) an audio-visual perception task using the same items as in the imagery tasks. In the famous-faces imagery task, participants had to decide which celebrity had a more round or oval face. Participants heard a word indicating a particular imagery category (e.g., “face”), followed by 2 names, designating the items the participant is required to imagine (e.g. “E.Macron”, “F.Hollande”, stimulus onset asynchrony = 4s). They were instructed to generate and maintain mental images as vivid as possible for 4 s per item. Eight seconds after the onset of the second item, they heard an attribute word (e.g. “round”). They then pressed one of two buttons accordingly, indicating which of the items is most closely associated with the attribute (e.g. who is associated with the attribute “round” face). In the color imagery task, participants had to decide which fruit or vegetable had a darker or lighter color. In the non-imagery abstract words control task, participants had to decide which of two abstract words (e.g., “routine”, “convention”) was semantically closer to a third word (e.g. “society”). In the perception task, the same items used for the imagery tasks were presented with both the auditory word and a visual image. In the abstract-word and the perception tasks, participants rated their confidence on a 4-level Likert scale, instead of rating vividness.

Each participant was scanned for 120 minutes at maximum and finished 5 runs. The first 3 runs consisted of 90 imagery-only trials (30 trials per run, each including 6 trials per category). The last 2 runs consisted of 36 trials each, with 3 trial types presented in a specific order (6 abstract word control trials, 15 imagery trials and 15 auditory-visual perception trials per run, each including 3 trials per category). Notably, the last two runs contained identical items in the imagery trials and perceptual trials, and participants only saw the perceptual items after the imagery trials. Trials within each trial type were randomized across categories.

### Dataset 2: passive reading

In this dataset^7^ using block design, 21 participants (mean age=23.4, 10 female) with varied language dominance in English and French passively viewed 6-letter strings presented at a fast pace (stimulus presented for 150 ms, followed by a fixation of 200 ms), while detecting catch trials with hashtags (“######”) by pressing a button. The letter strings consisted of 14 conditions (4 conditions each for letters, bigrams, quadrigrams, 2 conditions for real words in English and French), where the word-component frequency was separately modulated in English and French. Each condition was presented in short blocks, repeated 15 times. This main experiment lasted for 39.9 minutes, split into 3 runs. An independent object category localizer run was acquired, with conditions including faces, bodies, English words, French words, numbers, false fonts, houses, tools, checkerboards, each repeated 5 times. The localizer experiment lasted 9.2 minutes. Within each experiment, all blocks were randomized. The whole scanning session lasted around 90 minutes. See detailed description of the experimental design in Ref.^7^.

### Preprocessing

We processed the anatomical and functional MRI data with BrainVoyager (Versions 22.0.2.4572 and 22.4.4.5188, Brain Innovation, Maastricht, The Netherlands, https://www.brainvoyager.com/), as previously reported^7,22^. All analyses here are conducted with BrainVoyager, MATLAB (version R2018b), NeuroElf 1.1 toolbox (http://neuroelf.net/) implemented in MATLAB, and JASP 0.16.2 (https://jasp-stats.org/). GLMs were computed with predictors including each single condition, the response period, the button presses, and the 6 parameters for head motion (z-normalized). Functional voxel size is 1.2mm isotropic.

To evaluate data quality, particularly in the OFC, which is susceptible to signal dropout, we computed temporal signal-to-noise ratio (tSNR) maps for each participant and run, from the preprocessed, unsmoothed functional data. For each voxel, tSNR was defined as the temporal mean divided by the temporal standard deviation of the time series. tSNR was then averaged across voxels within each individual ROI, separately for each run (Dataset 1: runs 4 and 5, 439 volumes each; Dataset 2: runs 1–3, 399 volumes each). Differences between runs and between aphantasic and typical imagers were tested with paired and independent-samples t-tests, respectively. Results are reported in Fig. S11.

### Analyses for Dataset 1

We performed various analyses by integrating data from multiple runs:

#### Patch Identification

We used perceptual trials from runs 4 and 5 (6 trials per category) to identify face and color patches, as regions of interests (ROIs) in subsequent analyses.

#### Activation Amplitude, Category Selectivity, and Functional Connectivity

We used runs 4 and 5, where participants had identical items and number of trials for imagery and perception, with control trials using abstract words for functional connectivity. Note that ROIs were defined exclusively from perception trials (runs 4 and 5). All imagery analyses were therefore performed on data independent of ROI definition. For perception, we verified that the OFC selectivity and activation amplitude were preserved with cross-validated estimates, in which ROIs were defined on one run and beta values extracted from the other (Fig. S5B, C).

#### Neural representational content

We examined multi-voxel patterns for individual cortical patches. For the perceptual similarity matrix, we extracted beta values from perceptual trials in runs 4-5 (6 items per category, 6x6 matrices). For imagery similarity analysis, we extracted beta values from imagery trials in runs 1-3 (18 imagery trials per category, 18x18 matrices) to enhance the accuracy of estimation. To assess the representational overlap between imagery and perception for the exact same items, beta values were extracted from imagery and perceptual trials in runs 4-5. See the corresponding sections below for detailed methods.

### Identification of face- and color-preferring patches by univariate analysis

We performed general linear models (GLMs) on fMRI data of run 4 and 5, and used the contrast of faces or colors versus remaining categories in perceptual trials to identify face- or color-preferring patches as ROIs. Whole-brain voxelwise correction for multiple comparisons (e.g., FDR) is overly strict for small, individually variable patches defined from a limited number of trials, and would mainly increase false negatives in regions with weaker signals such as the OFC. We therefore used an uncorrected threshold (voxelwise p<0.001 uncorrected, cluster size thresholded at 12 voxels) for ROI definition and assessed the reliability of the OFC patches by independent cross-run replication (see Test of replicability of OFC patches). Because gradient-echo EPI at 7T is sensitive to signals from large draining veins, candidate patches that were likely of venous origin, e.g., clusters located on the pial surface or overlapping with veins visible on the anatomical or mean functional images, were excluded during ROI definition.

The VOTC face- and color-preferring patches were labeled according to conventions in the literature^9,29,79^. From posterior to anterior, face patches included: occipital face area (OFA), fusiform face area 1 (FFA1), fusiform face area 2 (FFA2), anterior face patch (AFP); color patches included: posterior color (PC), central color (CC), anterior color (AC) patches. Outside the VOTC, in bilateral OFC, we identified for faces (OFC-f) and for colors (OFC-c). Notably, with just 6 perceptual trials (each 24 seconds) per category, the OFC face and color patches could be identified in 18/20 and 19/20 individuals, respectively. For three participants without OFC patches in perceptual trials (Aph 01 and Aph 05 lacked face patches, Typ 04 lacked color patches), the OFC ROIs were defined by 18 imagery trials from independent data (run 1-3) with the same category-specific contrasts. For these participants, imagery analyses used runs 4–5 and thus remained independent of ROI definition. To test the medial-lateral differences between the face and color patches in both OFC and VOTC, and in both typical and aphantasic groups, we computed, for each participant and patch, the mean Talairach coordinates of all its voxels pooled across both hemispheres, using absolute values for the x coordinates. Group-average coordinates reported in the Results are the means of these individual coordinates across participants, such that each participant contributed equally regardless of patch size. For subsequent analyses, we created one ROI per area by combining the bilateral patches for each individual. To determine the activity profiles per patch, we extracted the beta values (% signal changes) per condition from run 4 and 5, where participants had identical items and number of trials for both imagery and perception.

### Test of replicability of OFC patches

To test whether the OFC patches replicate in independent data, the perception trials were split into ROI-definition and test datasets. In Dataset 1, OFC face (OFC-f) and color (OFC-c) patches were defined on one run (run 4 or run 5) and tested on the other, independent run with the same contrast: for OFC-f, face vs. the other categories (shape, color, word and map); for OFC-c, color vs. the other categories (shape, word, face and map). In the test run, the contrast was estimated for each ROI with a ROI-level GLM. In Dataset 2, OFC word patches were defined on two runs (word-similarity effect, p < 0.05) and tested on the remaining run (leave-one-run-out). One participant completed only two runs and contributed two folds. All 21 participants were included in this analysis, regardless of whether OFC word patches were identified in the full data. In the test run, the word-similarity effect was quantified as the Spearman correlation (ρ) between the ROI response and word similarity across the 14 conditions.

To generalize the individual-level results to the population, we used Bayesian prevalence inference^80^. For each participant and cross-validation fold, the test-run effect was classified as positive if it was in the predicted direction (t > 0 for face/color patches; ρ > 0 for word patches) and significant at p < 0.05 (two-sided, uncorrected). From the number of positive participants k out of n, we estimated the population prevalence γ, i.e., the proportion of the population that would show a true effect, under a uniform prior. We report the maximum a posteriori (MAP) estimate and the 96% highest posterior density intervals (HPDI). Each fold was analyzed separately. Differences in prevalence between aphantasic and typical imagers were assessed from 10^6^ posterior samples per group. Prevalence analysis used the MATLAB implementation of ref.^80^.

### Bayesian statistical tests and Bayes Factors

Statistical tests were performed using JASP 0.16.2 (https://jasp-stats.org/) with the JASP default priors, or using BayesFactor 0.9.12 implemented in R 4.6.1. A commonly accepted convention is that Bayes factors (BF10 or BFs) between 3 and 10 indicate moderate evidence in favor of the model in the numerator (H1); BFs between 10 and 30 indicate strong evidence; BFs larger than 30 indicate very strong evidence. The inverse of these cut-offs values provides moderate (<0.33), strong (<0.10), or very strong evidence (<0.03) for the model in the denominator (H0), i.e. the null hypothesis.

### Temporal profile analysis

We extracted the event-related average time courses across voxels in each ROI. The time courses were normalized by the averaged activity across TRs -2 to 0. We performed a three-way ANOVA to identify group differences across patches and time points (Group x Patch x TR [0 to 11], trials end at TR 12), separately for imagery and perception conditions. We performed three-way ANOVAs with the factor of Group x

Patch x TR in each condition. For both face perception and face imagery, there were three-way interactions and the group effect interactions with TR. We then unpacked the group effect for each time point in each of the ROIs and tested the group difference with time-resolved independent t-tests. Results were Bonferroni-corrected across time points with corrected alpha = 0.05, for each ROI. There were group differences for both color perception and color imagery, and these effects didn’t interact with other factors.

Detailed statistics for ANOVAs are reported in Fig. S4.

### Face- and color-selectivity index

For the face- and color-preferring patches, we used the activation amplitudes (single-condition beta values) to calculate the selectivity index as follows: Selectivity index (SI) for the preferred category = (4 * preferred category – other categories) / (4 * preferred category + other categories)

To avoid deactivation in the fMRI signal leading to an overestimation of selectivity, negative amplitudes were set to zero before computing the SI^31^. This adjustment resulted in SI values ranging from -1 to 1. A positive SI indicates category selectivity; a SI of 0 indicates similar amplitude across all conditions; a negative SI indicates that on average the region is activated more strongly by the other conditions.

### Behavioral multiple-arrangement tasks

After the fMRI scanning session, participants performed multiple-arrangement tasks^81^ on Meadows Research platform (https://meadows-research.com/). We sought to determine their subjective judgments on semantic or visual similarity of faces and colors. Participants were asked to drag and drop the items within the circle with a mouse, a pair at a time, and place items close to each other according to the considered similarity. For each category, participants arranged the same items in separate tasks, either by semantic features (cued by words), or perceived features (cued by photos). Specifically, for faces, semantic feature representational dissimilarity matrices (RDMs) derived from written names of the celebrities arranged according to their social identities, i.e. professions; the visual feature RDMs derived from photos arranged according to the facial shapes (round or elongated). For object colors, the semantic RDMs were derived from food items arranged by type (vegetables, tropical fruits, and prepared food); the visual feature RDMs derived from food items arranged from their visual color appearances. To map the neural RDMs separately for imagery and perception, participants arranged 6 items from perceptual trials regarding semantic and visual features (6x6), then arranged 18 items from imagery trials regarding semantic and visual features (18x18), resulting in 4 behavioral RDMs for each category. RDMs were calculated using Euclidean distance from Meadows Research. The resulting behavioral arrangements were visualized with multidimensional scaling^82^ from the RDMs from each group. Figure S6 displays exemplar arrangements of 18 items from imagery trials. In the same session, we also collected similarity judgments on object shapes, words and French maps with the items from the fMRI tasks of the Dataset 1, and for each category, participants also arranged items by relevant visual features using word cues. Those data were not used in the current analysis.

### Statistical tests on behavioral RDMs

We first tested the level of inter-individual variability. We calculated leave-one-out squared difference between each individual and within group means. The test statistic was the sum of the upper triangular elements of the difference matrix with a dimension of 153 (for 18 items), resulting in one value for each participant and for each RDM. We then performed independent t-tests to compare the two groups. No group difference was observed in any of the RDMs (all ps>0.12). We then performed nonparametric permutation tests to evaluate group differences on each behavioral RDMs. For each permutation, participant labels were randomly shuffled while maintaining the original group sizes. The squared sum of mean differences was calculated for each permuted dataset. This procedure was repeated for 1000 permutations. The observed test statistic from the actual data was compared to the distribution of the permuted statistics. A p-value was calculated as the proportion of permuted statistics greater than or equal to the observed statistic, with a cut-off p-value of 0.05 indicating significance.

### Neural RDMs

For each ROI, we first obtained voxel-wise average % signal change from the 4-5th repetition times (TRs, normalized by the TRs -2 to 0 as the baseline), which corresponded to the peak activity of the first item, with minimal influence from the second item in each trial. We then generated the neural RDM across stimulus pairs with Pearson correlation. Furthermore, to directly assess the similarity of neural representations between imagery and perception (representational overlap) within each patch, we correlated RDMs (6x6) generated from the run 4-5 where participants imagined or perceived the same 6 items. We then examined the amount of overlap by Bayesian one-sample t-tests against zero for each group and Bayesian independent t-test between groups on each patch.

### Linear regression with RDMs

In each patch, separately for faces and colors, to examine the presence of semantic or visual-feature representational content during perception and imagery, we fitted a linear regression model, using two behavioral RDMs (semantic, visual) as regressors, and the neural RDMs as dependent variables. The behavioral RDMs have been squared before model fittings.

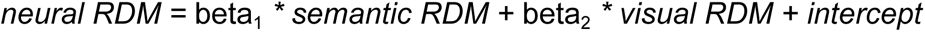

We tested the beta weights per patch using Bayesian one-sample t-tests against zero to determine the presence of representational contents. No comparison was made between the semantic and visual beta values.

### Psychophysiological interactions (PPI) analysis

To investigate task-specific functional connectivity for each individual ROI, we conducted PPI analysis with trials in run 4-5. We built each PPI design matrix by 1) generating a task contrast predictor between two conditions (Condition A - B), balanced by a regressor of the sum of those two conditions (Condition A + B), both convolved with the hemodynamic response function (HRF); 2) extracting a demeaned time course from the seed ROI; 3) generating an interaction predictor as an element-by-element product of the HRF-convolved task contrast predictor and the seed time-course predictor; 4) adding the 6 parameters of participant’s z-scored head motion as confounding predictors. Specifically for 1), to investigate category-specific connectivity, we built the task contrast predictor by subtracting the Abstract word control condition from the single-category condition under investigation, e.g. face imagery, while keeping the other imagery / perceptual category conditions unchanged. We separated PPI design matrices for each seed ROI and each contrast. The seed ROIs included OFA, FFA1, FFA2, AFP, OFC-f for face imagery and perception; and PC, CC, AC, OFC-c for color perception and imagery. We then computed GLMs on individual unsmoothed fMRI datasets in BrainVoyager, resulting in whole-brain beta maps of each task contrast. For each seed ROI, its connectivity to every other ROI was averaged across voxels for the resulting beta values per ROI. To examine the bottom-up influence from early visual cortices, we also included V1 as a seed, which was manually delimited anatomically in each individual brain around the retro calcarine sulcus. A complementary group-level whole-brain analysis (smoothed data, FWHM = 6 mm) is described in Supplementary Materials (Fig. S7).

### Analyses for dataset 2

#### Definition of OFC word-preferring patches

We used the predictor for the word-similarity effect (across 14 conditions, [1 2 2 3 4 5 5 6 7 8 8 9 10 10], see Ref.^7^, and performed Spearman correlation across 14 conditions on the single-condition beta values per voxel. This predictor assumes increasing word-component frequency across letters, bigrams, quadrigrams, and real words, but does not assume a frequency difference of the same word component between English and French. The resulting Spearman Rho maps were thresholded at p<0.005 uncorrected (functional cluster size > 4) to mitigate false negatives. False positives were controlled by the independent cross-run replication test (Fig. S10). Since OFC word-preferring patches were found in 10/21 participants, all subsequent descriptive statistics and statistical analyses were performed on those 10 participants. To assess word selectivity in the independent object category localizer, ROI GLMs of the averaged time course per patch was computed in BrainVoyager (Figure 5A black arrowheads indicating significant patches), and the activation amplitudes (beta values) averaged across all OFC patches per participant were extracted (Figure 5B and Figure S8). The word SI was computed from the Dataset localizer betas, with the average of English and French words as the preferred category and faces, bodies, houses and tools as the other categories. As in Dataset 1, negative amplitudes were set to zero before computing the SI.

### Definition of VOTC word- and face-preferring patches

As in Zhan et al.^7^, word-preferring patches in the VOTC were defined with the object category localizer (contrast: English and French words > faces, bodies, houses, tools; p < 0.001 uncorrected; cluster size > 4 functional voxels). Candidate patches that were likely of venous origin, e.g., clusters located on the pial surface or overlapping with veins visible on the anatomical or mean functional images, were excluded during ROI definition. Face-preferring patches were also defined (contrast: faces > bodies, houses, tools, average of English and French words), as a reference for their medial-lateral layouts relative to the word patches in the VOTC. The anatomical locations of all patches were manually labeled (ROI data are publicly available on https://osf.io/96syx/?view_only=88b55034027042fbb4f118f398fc5706). All patches around the fusiform gyrus were merged into a single ROI per hemisphere.

### Comparison of the medial-lateral layouts between word and face patches

For both OFC and VOTC patches, within each participant, the absolute value of the average Talairach x coordinates across voxels were computed within each hemisphere, and were then averaged across hemispheres. The resulting x coordinates of the VOTC word and face patches were compared within the 10 participants with a paired-sample t-test. Since the localizer in Dataset 2 did not reveal reliable OFC face patches, the x coordinates of the OFC word patches were compared to the x coordinates of the OFC face patches in Dataset 1, with a Wilcoxon rank sum test for independent samples (10 versus 20 participants).(END)

