## Supplementary material (v202610) for "Category-preferring patches in human orbitofrontal cortex"

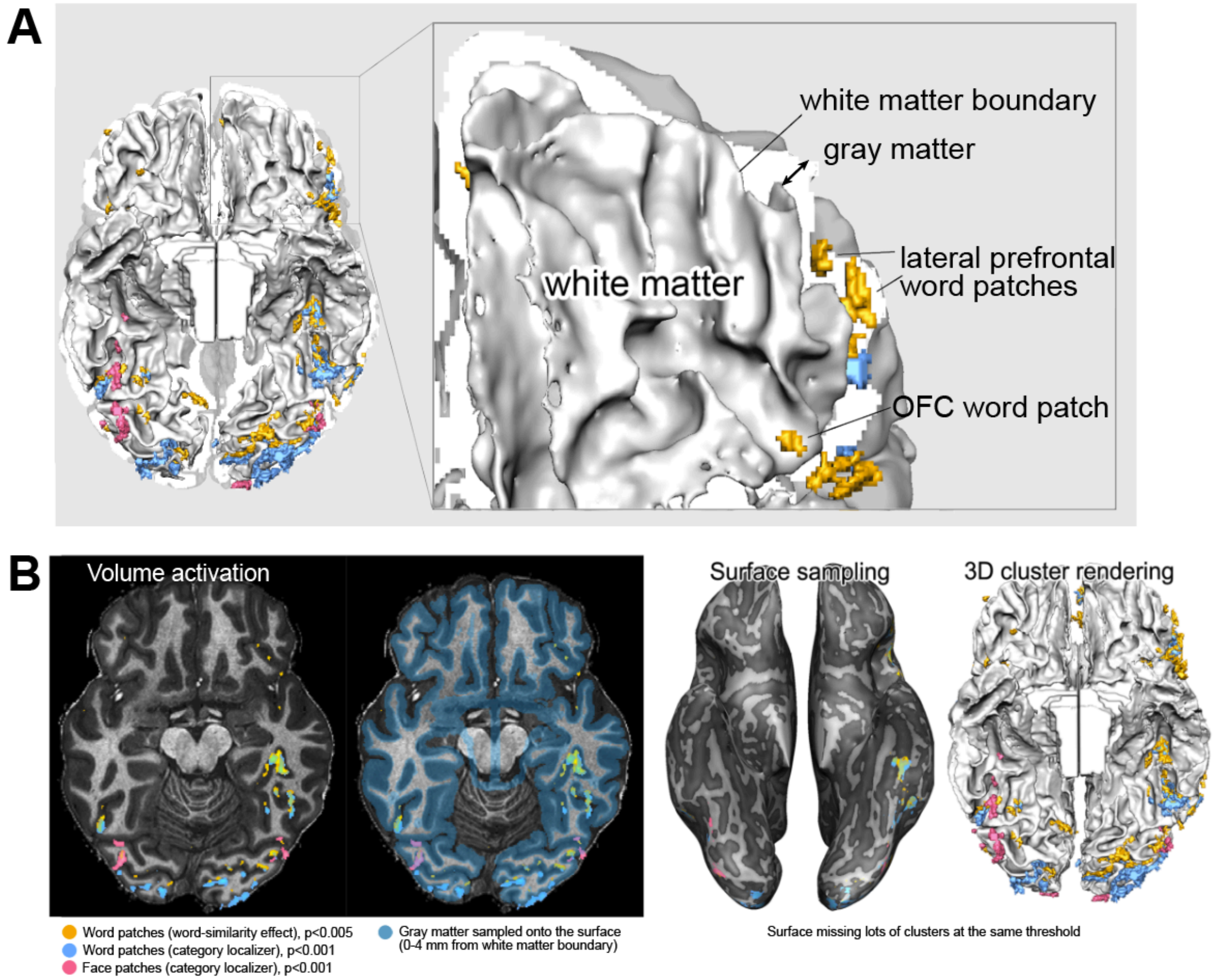


**Fig. S1 Visualization of functional patches relative to the cortical surface.**

**A.** Functional maps were computed in volume space and displayed as 3D-rendered clusters on the gray-white matter boundary. Data from one representative participant (SE03) of Dataset 2. The gray matter is rendered in white color to show the cortical thickness. Because the patches are located within the gray matter, i.e., outside the white-matter surface, they appear outside or partially occluded by the rendered surface depending on the viewing angle. Left: ventral view of the whole brain. Right: magnified view of the orbitofrontal cortex, showing an OFC word patch, and lateral prefrontal word patches (not analyzed in the paper) lying in the gray matter outside the gray-white matter boundary.

**B.** Comparison of visualization methods for the same functional maps and thresholds. From left to right: activation in volume space overlaid on a transversal slice of the anatomical image; the range of gray-matter sampling depth is 0-4 mm for surface projection (blue shading); the maps sampled onto the inflated cortical surface; and the 3D cluster rendering used in Fig. 1A and Fig. 5A. Some clusters present in the volume space far from the gray-white matter boundary failed to be captured by surface projection at the same thresholds, but were preserved in the 3D rendering.

### Test of replicability of OFC face and color patches

Perceptual trials were split into ROI-definition and test datasets. OFC face (OFC-f) and color (OFC-c) patches were defined on one run (run 4 or 5) and tested on the other run with the same contrast (OFC-f: face vs. shape, color, word and map; OFC-c: color vs. shape, word, face and map) using a ROI-level GLM computed on the averaged time course per ROI. To generalize individual-level results to the population, we used Bayesian prevalence inference. For each participant and fold, the test-run effect was classified as positive if $t > 0$ and $p < 0.05$ (two-sided, uncorrected). This test has a false-positive rate of at most 0.025; we conservatively set $\alpha = 0.05$ in the model. From the number of positive participants, we estimated the population prevalence under a uniform prior and reported the maximum a posteriori (MAP) estimate and the 96% highest posterior density intervals (HPDI). Each fold was analyzed separately. Differences between aphantasic and typical imagers were assessed from $10^6$ posterior samples per group.

OFC face and color patches defined in one run showed significant category preference in the independent test run (Fig. S2). At the group level, test-run t-values were significantly greater than zero for OFC-f (mean $t = 3.49$ and 3.76 for ROIs defined in run 4 and run 5; one-sample t-tests, $t(19) = 7.09$ and 6.28, both $p < 0.001$) and for OFC-c (mean $t = 3.18$ and 2.78; $t(19) = 8.03$ and 9.85, both $p < 0.001$). At the individual level, OFC-f patches were replicated in 16 of 20 participants in both directions, and OFC-c patches in 14 and 13 of 20 participants for ROIs defined in run 4 and run 5, respectively (Table S1). Bayesian prevalence analysis for population-level inference estimated that the majority of the population would show replicable OFC patches: MAP = 0.79 (96% HPDI [0.57, 0.93]) for OFC-f in both directions, and MAP = 0.68 ([0.45, 0.86]) and 0.63 ([0.40, 0.82]) for OFC-c. Prevalence did not differ between aphantasic and typical imagers: the 96% HPDIs of the difference included zero in all folds ($\gamma AP - \gamma HP$: MAP = −0.10 to 0.19; $P(\gamma AP > \gamma HP) = 0.33$–0.85). Together, these analyses show that the OFC patches replicate across independent runs in the majority of participants, providing converging evidence for a reliable population-level effect.

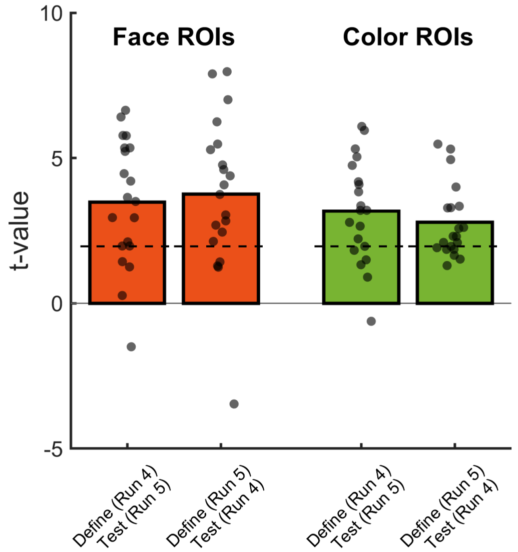


**Fig. S2 Cross-run validation analysis of OFC face and color patches.**
Bars represent the group-averaged t-values on test runs (n =20). Dots represent t-values in individual participants. Dashed lines indicate uncorrected within-participant significance levels ($p<0.05$, $t=1.96$).

**Table S1. Test of replicability of OFC face and color patches and prevalence summary**

| Group | ROI Type | Define (run) | Test (run) | First-level Prevalence | MAP | HPDI96 Lower | HPDI96 Upper |
|---|---|---|---|---|---|---|---|
| **All** | **color** | **4** | **5** | **14/20** | **0.68** | **0.45** | **0.86** |
| Aph | | | | 8/10 | 0.79 | 0.48 | 0.96 |
| Typ | | | | 6/10 | 0.58 | 0.27 | 0.84 |
| **All** | **color** | **5** | **4** | **13/20** | **0.63** | **0.40** | **0.82** |
| Aph | | | | 6/10 | 0.58 | 0.27 | 0.84 |
| Typ | | | | 7/10 | 0.68 | 0.37 | 0.91 |
| **All** | **face** | **4** | **5** | **16/20** | **0.79** | **0.57** | **0.93** |
| Aph | | | | 9/10 | 0.89 | 0.60 | 0.99 |
| Typ | | | | 7/10 | 0.68 | 0.37 | 0.91 |
| **All** | **face** | **5** | **4** | **16/20** | **0.79** | **0.57** | **0.93** |
| Aph | | | | 9/10 | 0.89 | 0.60 | 0.99 |
| Typ | | | | 7/10 | 0.68 | 0.37 | 0.91 |

MAP: maximum a posteriori estimate of population prevalence; HPDI: highest posterior density interval. Aph: aphantasic individuals; Typ: typical imagers.

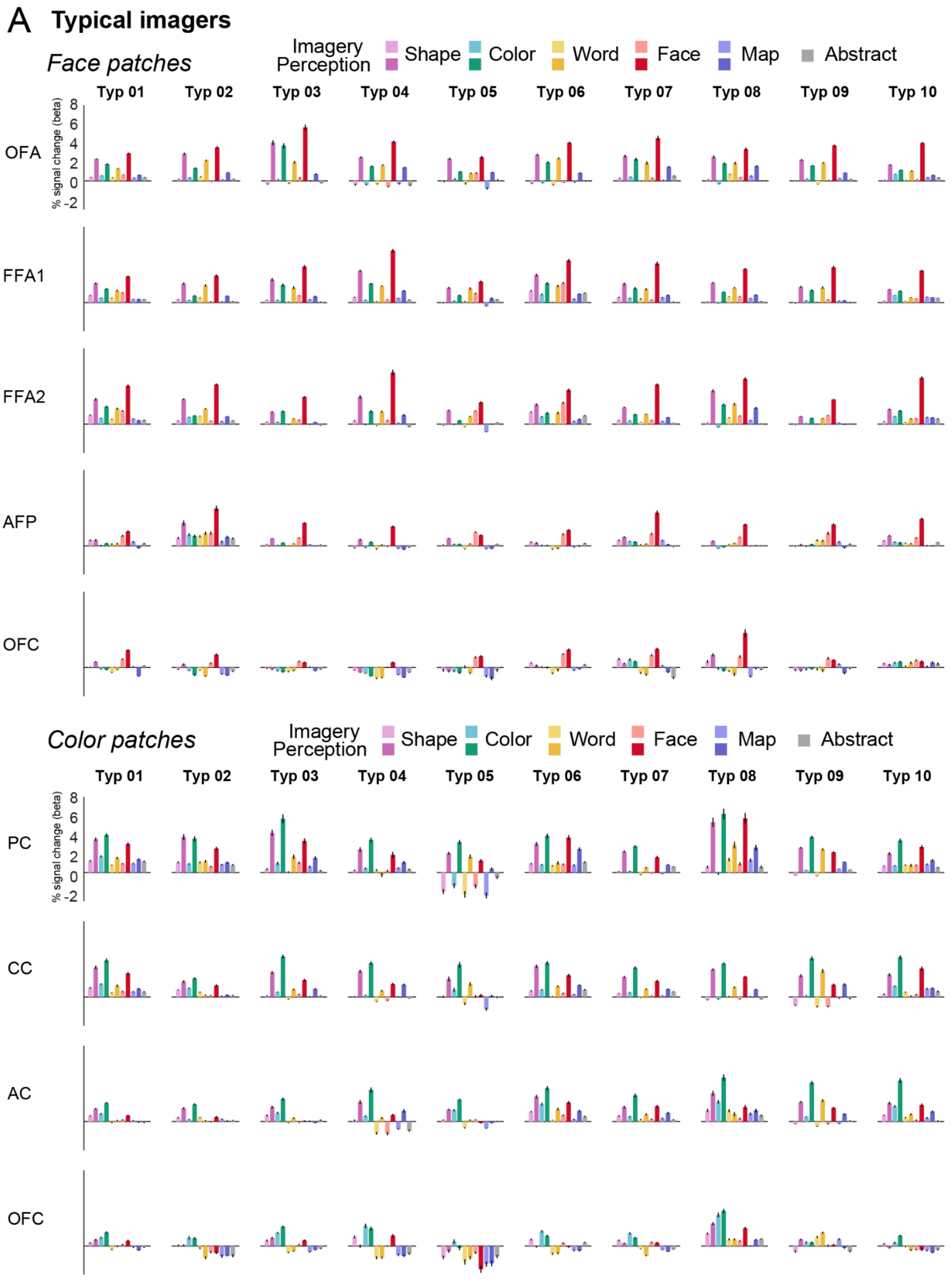


(continued on the next page)

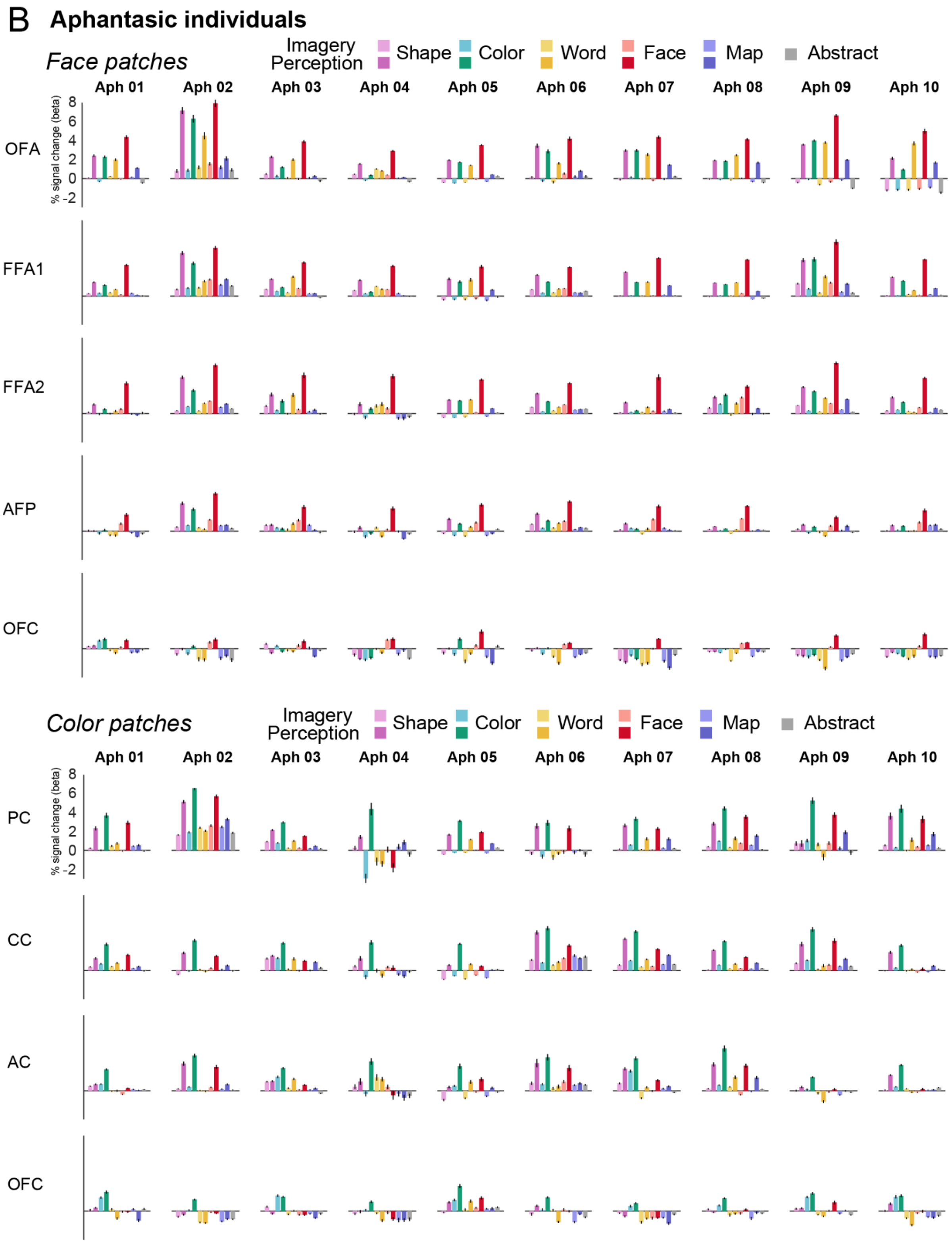


**Fig. S3 Individual activity profiles of face and color patches across tasks**
(A) Typical imagers. (B) Aphantasic individuals. The below-baseline activity for non-preferred categories in OFC patches was observed in individual participants.

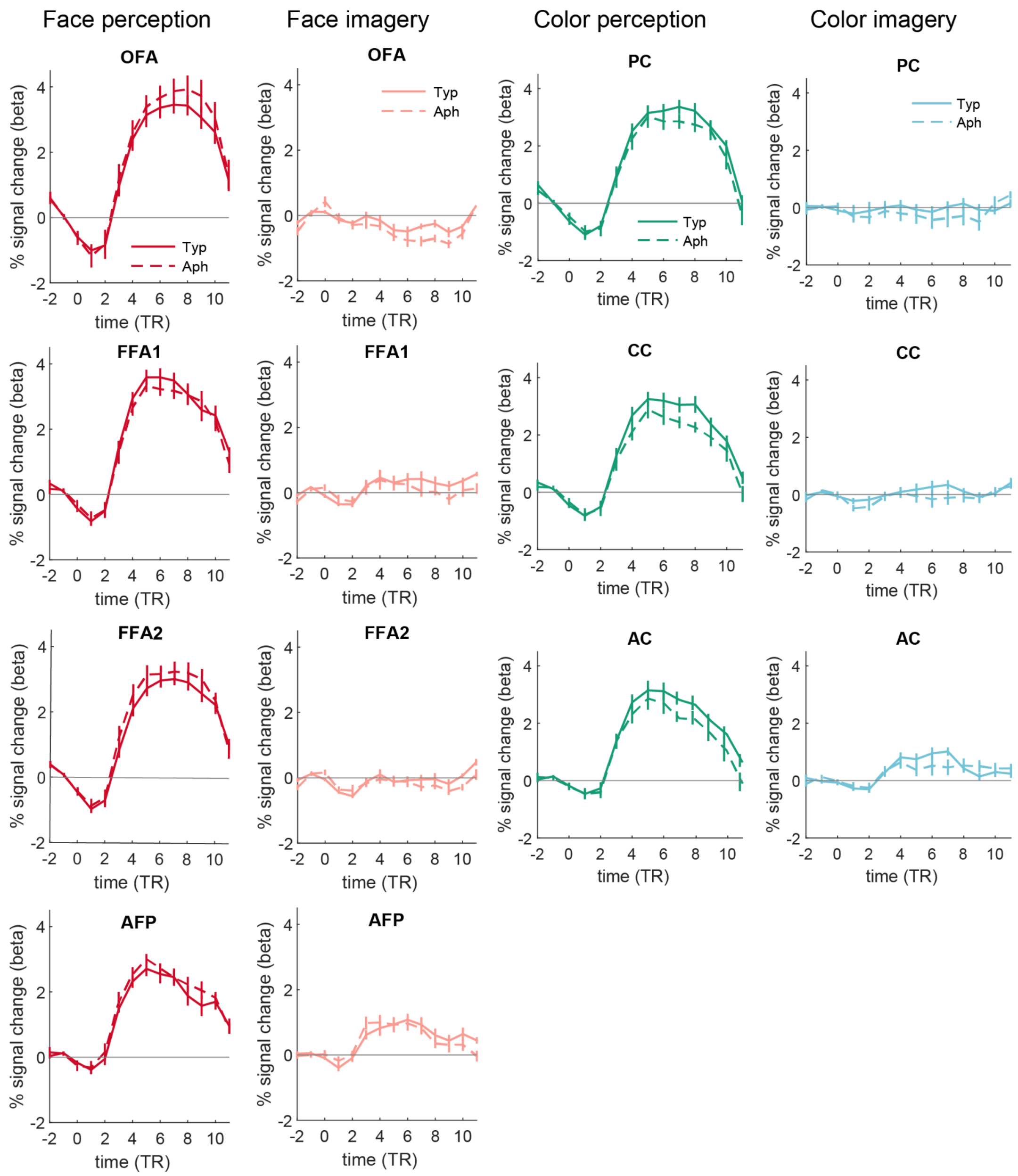


**Fig. S4 Temporal profiles of VOTC patches.**
We performed three-way ANOVAs for each condition with the factor of Group x Patch x TR. In face perception, there was a three-way interaction ($F = 2.67$, $p<0.001$, $\eta^2 p = 0.096$) although the main group effect was not significant ($F = 3.27$, $p = 0.07$, $\eta^2 p = 0.004$). The group effect interacted with TR ($F = 85.81$, $p < 0.001$, $\eta^2 p = 0.462$). There were main effect of patch ($F = 8.25$, $p < 0.001$, $\eta^2 p = 0.035$), TR ($F = 218.37$, $p < 0.001$, $\eta^2 p = 0.686$) and the patch x TR interaction ($F=7.82$, $p <0.001$, $\eta^2 p = 0.238$). In face imagery, there was a three-way interaction ($F = 1.69$, $p=0.007$, $\eta^2 p = 0.063$) without main group effect ($F = 0.009$, $p = 0.92$, $\eta^2 p < 0.001$). There was significant group x TR interaction ($F = 12.67$, $p <0.001$, $\eta^2 p = 0.112$). There were main effects of patch ($F = 171.43$, $p < 0.001$, $\eta^2 p = 0.432$) and TR ($F = 10.10$, $p < 0.001$, $\eta^2 p = 0.092$), and the patch x TR interaction ($F= 6.61$, $p<0.001$, $\eta^2 p = 0.209$). In color perception, there was a

main group effect ($F=23.32$, $p < 0.001$, $\eta^2p = 0.026$) with no interaction with other factors (all ps > 0.139). There were significant main effects of patch ($F = 12.56$, $p < 0.001$, $\eta^2p = 0.042$) and TR ($F = 184.89$, $p <0.001$, $\eta^2p = 0.702$), together with patch x TR interaction ($F = 6.47$, $p < 0.001$, $\eta^2p = 0.198$). In color imagery, there was a main group effect ($F=5.96$, $p=0.015$, $\eta^2p = 0.007$) without interaction with other effects. However, no time point showed a significant group difference in any VOTC patches after correction. There was also main effect of patch ($F = 107.69$, $p<0.001$, $\eta^2p = 0.272$), main effect of TR ($F = 11.77$, $p <0.001$, $\eta^2p = 0.130$) and significant patch x TR interaction ($F = 3.87$, $p< 0.001$, $\eta^2p = 0.129$). Other effects were not significant.

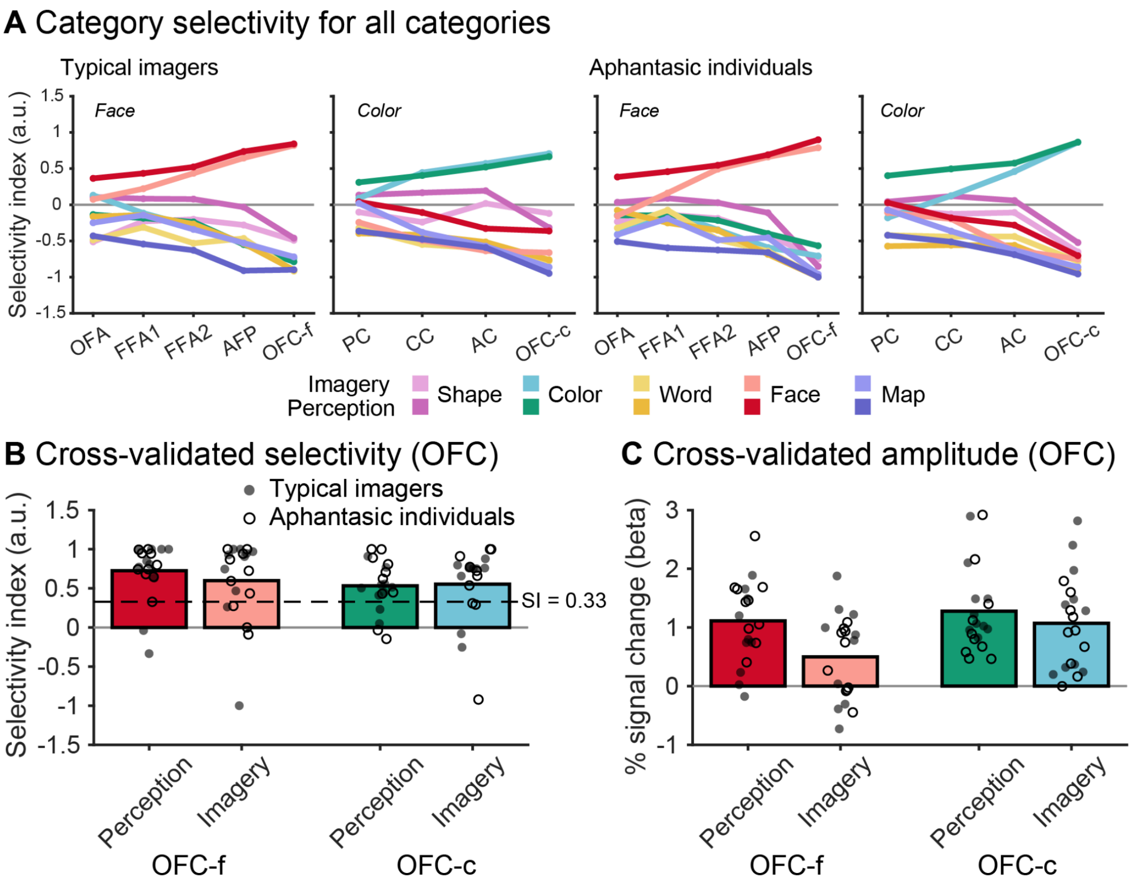


**Fig. S5 Category selectivity and cross-validated responses of OFC patches.** **(A)** Selectivity index (SI) for preferred and non-preferred stimuli in face- and color-specific patches of typical imagers and aphantasic individuals. Lines denote group averages; lighter colors: imagery; darker colors: perception. The OFC patches showed strong positive selectivity only for their preferred category, during both perception and imagery, and strongly negative selectivity for non-preferred categories. A negative SI indicates that on average the remaining categories have a higher activation amplitude. **(B, C)** Cross-validated SI (B) and activation amplitude for the preferred category (C) of OFC patches. ROIs were defined on one run, and SI and amplitude were computed from the other run (averaged across the two folds). Bars represent group averages ($n = 20$); filled and open dots represent typical imagers and aphantasic individuals, respectively. The dashed line in B indicates SI = 0.33 (activity for the preferred category twice that for the other categories). **(B)** The SI was above 0.33 in all conditions (one-sided one-sample t-tests; OFC-f perception: mean = 0.73, $t(19) = 4.92$, $p < 0.001$; OFC-f imagery: 0.60, $t(19) = 2.35$, $p = 0.015$; OFC-c perception: 0.53, $t(19) = 2.78$, $p = 0.006$; OFC-c imagery: 0.55, $t(19) = 2.09$, $p = 0.025$). **(C)** Activation amplitude was above baseline in all conditions (two-sided one-sample t-tests; OFC-f perception: 1.11 % signal change, $t(19) = 7.25$, $p < 0.001$; OFC-f imagery: 0.50, $t(19) = 3.14$, $p = 0.005$; OFC-c perception: 1.28, $t(19) = 7.89$, $p < 0.001$; OFC-c imagery: 1.07, $t(19) = 6.04$, $p < 0.001$). Neither SI nor amplitude differed between groups (independent samples t-tests, all ps > 0.10).

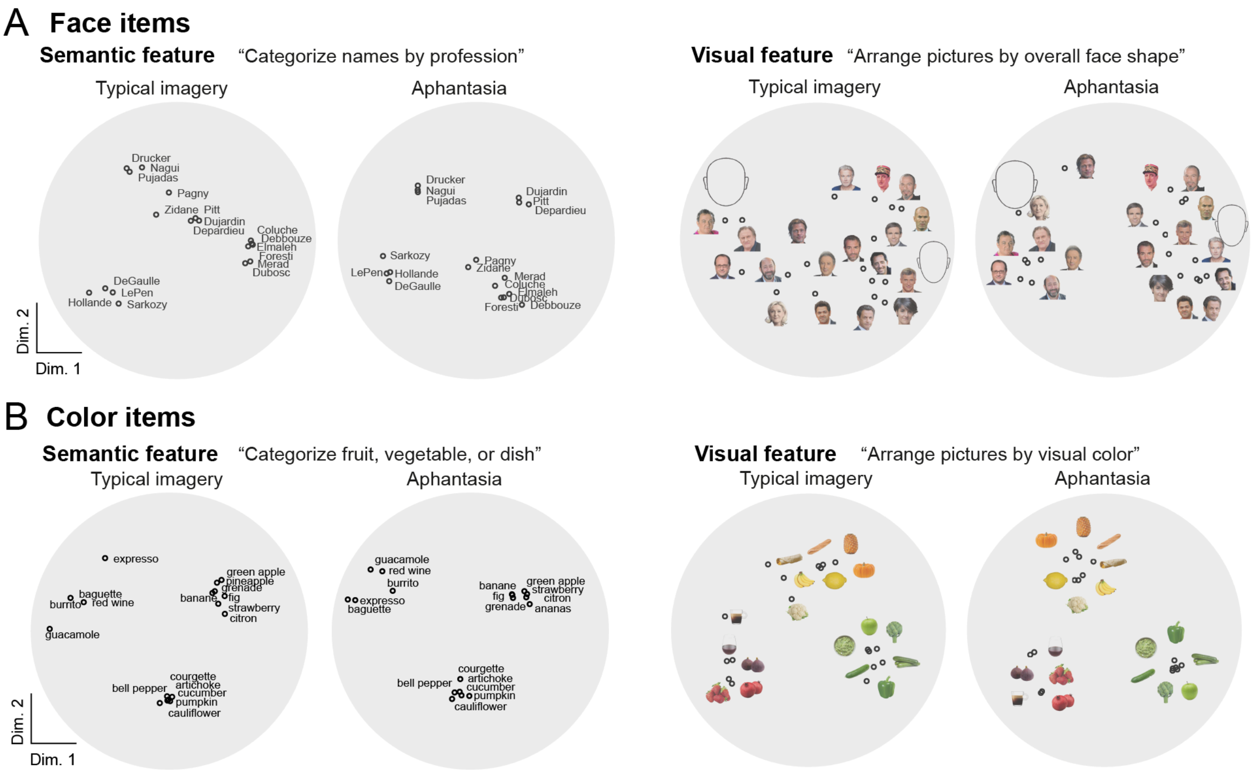


**Fig. S6 Behavioral similarity arrangements for items in the fMRI imagery trials.**

**A.** Visualization of face-item representations. Group averaged two-dimensional spaces were obtained through multidimensional scaling on individual arrangements of each group. In order to visually compare the spaces, we arbitrarily rotated the main axes of the dimensions while maintaining the relative position between the items within the space to align the groups. Dim.: dimension.

**B.** Visualization of color-item representations.

### Group-level functional connectivity of the OFC face and color patches

To characterize the whole-brain connectivity of the OFC patches, we performed seed-based psychophysiological interaction (PPI) analyses using the individually defined OFC-f and OFC-c patches (bilateral patches merged) as seeds. PPI contrasts were face perception vs. abstract words (OFC-f seed) and color perception vs. abstract words (OFC-c seed), constructed as described in Methods. For this group-level analysis only (N = 20), functional data were spatially smoothed (Gaussian kernel, FWHM = 6 mm) to account for inter-individual anatomical variability and allow overlap across participants; all individual-level analyses in this study were performed on unsmoothed data.

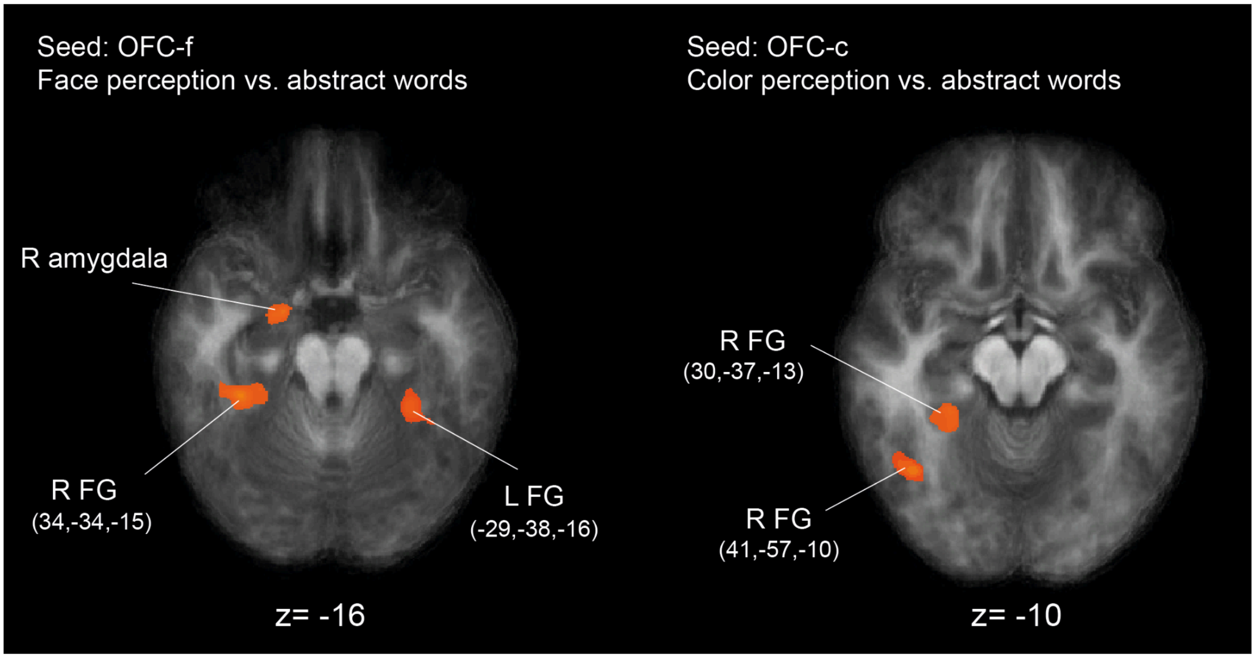


**Fig. S7 Whole-brain functional connectivity of OFC face and color patches.** Group-level (N = 20) PPI maps using OFC-f (left; face perception vs. abstract words) and OFC-c (right; color perception vs. abstract words) as seeds. Voxelwise $p < 0.005$, cluster-size corrected (Monte Carlo simulation, 5000 iterations, minimum cluster size = 100 voxels). Data were smoothed (FWHM = 6 mm) for group-level analysis. R/L: right/left; FG: fusiform gyrus. During face perception, the OFC-f seed was functionally coupled with the bilateral fusiform gyrus (FG; right: x = 34, y = −34, z = −15; left: x = −29, y = −38, z = −16) and the right amygdala. The FG peaks lay between the anterior face patch (AFP; group-average x = 36, y = −23, z = −16) and FFA2 (x = 37, y = −47, z = −15) identified at the individual level. During color perception, the OFC-c seed was coupled with two sites in the right FG. The anterior peak (x = 30, y = −37, z = −13) closely matched the anterior color patch (AC; x = 29, y = −34, z = −15), and the posterior peak (x = 41, y = −57, z = −10) was located at the level of the central color patch (CC; x = 30, y = −51, z = −12), but more laterally. Thus, at the whole-brain level,

OFC patches were preferentially coupled with anterior VOTC regions corresponding to the category-specific patches.

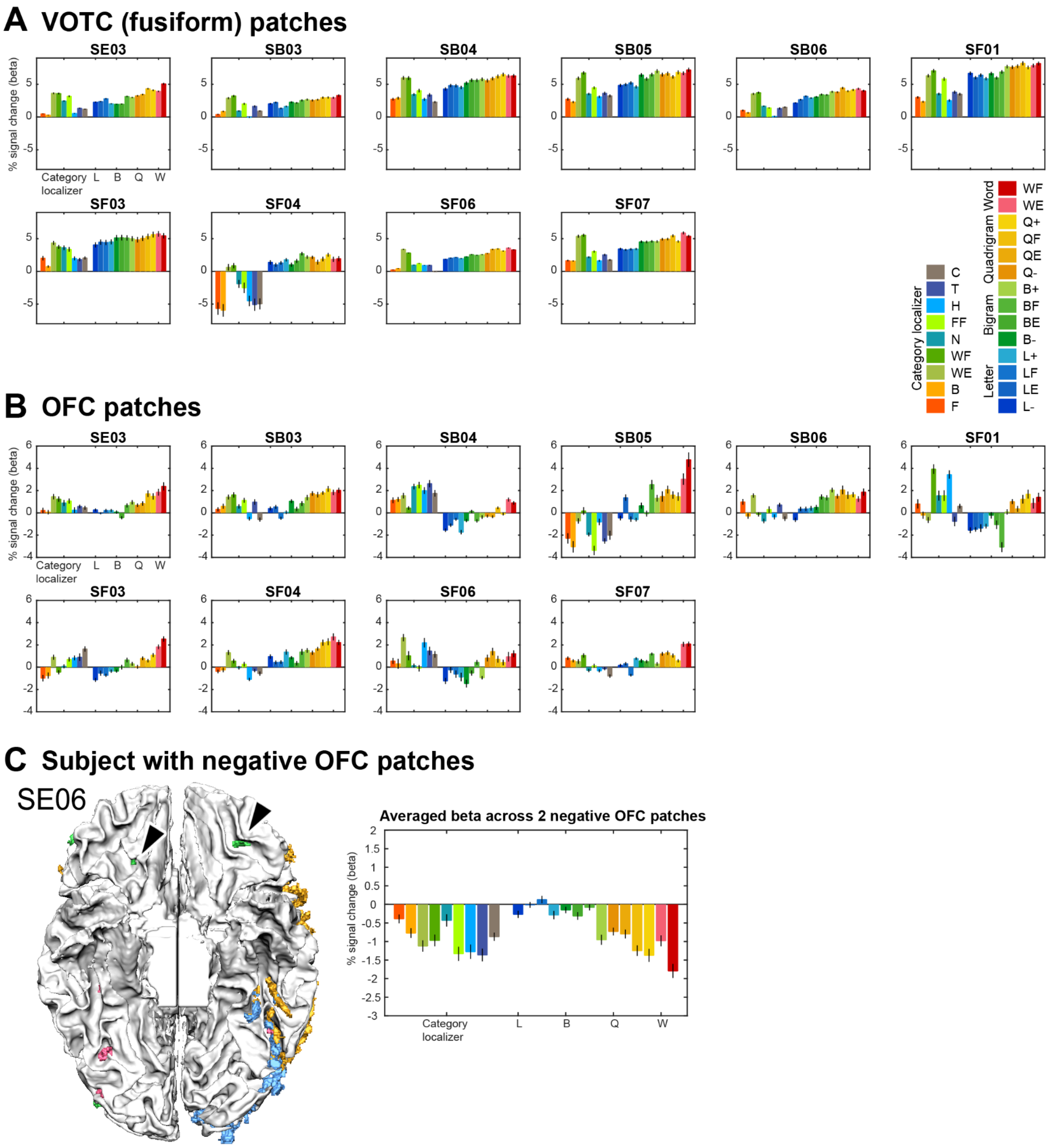


**Fig. S8 Individual-subject activity profiles of word patches, in Dataset 2.**
Within each panel, the left-side 9 conditions are from the object category localizer, the right-side 14 conditions are from the main experiment. Note that the VOTC ROIs in (A) are defined by the localizer data, and the OFC ROIs in (B) are defined by the main experiment data. Patches within VOTC or OFC were respectively merged into a single ROI per participant. Error bars denote the SEM across voxels within each individual participant. Abbreviations for conditions in the category localizer: F: faces, B: bodies, WE: English words, WF: French words, N: numbers, FF: false fonts, H: houses, T: tools, C: checkerboards; in the main experiment: L: letters, B: bigrams, Q: quadrigrams, W: words, E or F: word component frequency high in only English or only French, - or +: word component frequency low or high in both English and French. **(C)** Two negative OFC word patches in one subject, and the averaged beta values.

**Table S2. Ability to observe OFC word and face patches in Dataset 2, at different thresholds in individual participants.**

| Participant | OFC word (similarity slope) | | | OFC word (localizer) | | | OFC face (localizer) | | |
|---|---|---|---|---|---|---|---|---|---|
| | 0.005 | 0.01 | 0.05 | 0.005 | 0.01 | 0.05 | 0.005 | 0.01 | 0.05 |
| SE03 | + | + | + | - | - | + | - | - | + |
| SB03 | + | + | + | + | + | + | - | - | + |
| SB04 | + | + | + | - | - | - | - | - | - |
| SB05 | + | + | + | - | - | + | - | - | - |
| SB06 | + | + | + | + | + | + | + | + | + |
| SF01 | + | + | + | - | - | + | - | - | - |
| SF03 | + | + | + | - | - | - | - | - | - |
| SF04 | + | + | + | - | + | + | - | - | - |
| SF06 | + | + | + | - | - | - | - | - | - |
| SF07 | + | + | + | - | - | + | - | - | + |
| SE06 | +n | +n | +n | - | - | - | - | - | - |
| SF05 | - | +n | +n | - | - | - | - | - | - |
| SE01 | - | + | + | - | - | - | + | + | + |
| SB01 | - | + | + | - | - | - | - | - | + |
| SB02 | - | + | + | - | - | - | - | - | + |
| SE04 | - | + | + | - | - | - | - | - | + |
| SE07 | - | - | + | - | - | + | - | - | + |
| SB07 | - | - | + | - | - | + | - | - | + |
| SF02 | - | - | +n | - | - | - | - | - | - |
| SE02 | - | - | - | - | - | - | + | + | + |
| SE05 | - | - | - | - | - | - | - | + | + |

The patches are defined with the word similarity slope, the localizer word contrast, and the localizer face contrast, under 3 different thresholds: $p<0.005$, $p<0.01$, $p<0.05$. Cluster size > 4 functional voxels. +: OFC patch observable; -: OFC patch not observable. n: patch showing a negative word similarity slope, where stimuli less similar to real words elicited higher activity.

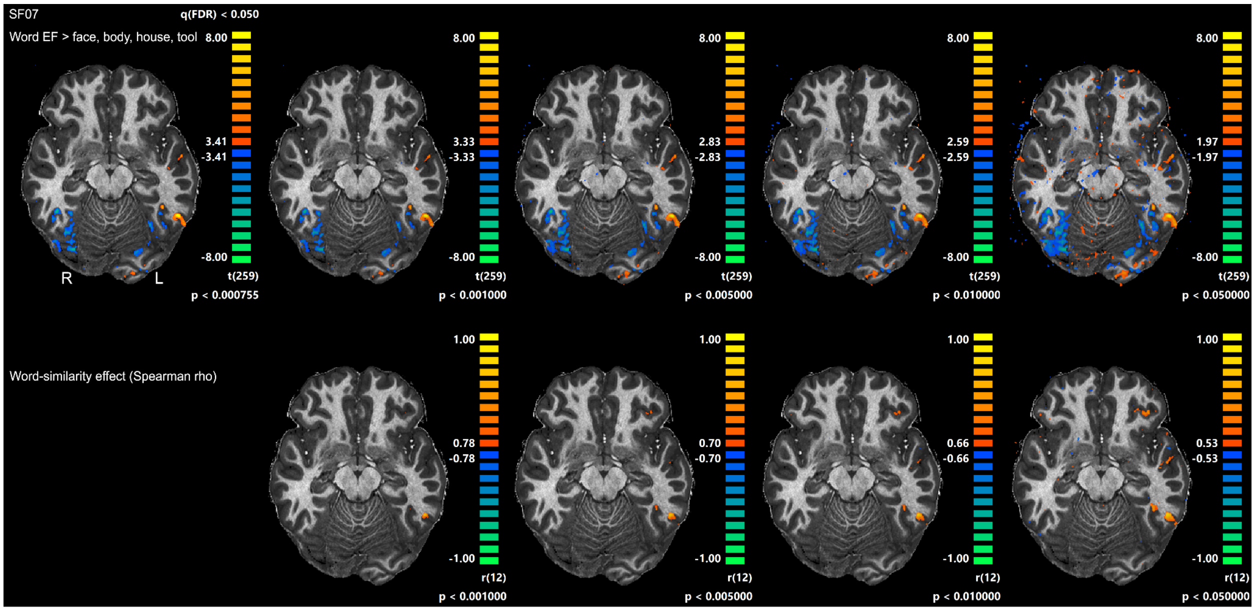


**Fig. S9 OFC and fusiform word patches in a representative subject** (SF07), under different thresholds (FDR, p<0.001, p<0.005, p<0.01, p<0.05). The patch sizes did not change dramatically in more lenient thresholds.

### Test of replicability of OFC word patches

OFC word patches were defined on two runs ($p < 0.05$) *in all 21 participants*, and tested on the remaining run (leave-one-run-out). In the test run, the word-similarity effect was quantified as the Spearman correlation (ρ) between ROI response and word similarity across 14 conditions.

The word-similarity effect in the test runs was significant at the group level (mean ρ = 0.40, 0.34 and 0.35 for test runs 1–3; one-sample t-tests on Fisher-z-transformed ρ, t = 4.22, 3.90 and 4.39, all ps < 0.001). It was significant in 12/21, 9/21 and 8/20 participants (ρ > 0 and $p < 0.05$; two-sided, uncorrected), see estimated prevalence in Table S3.

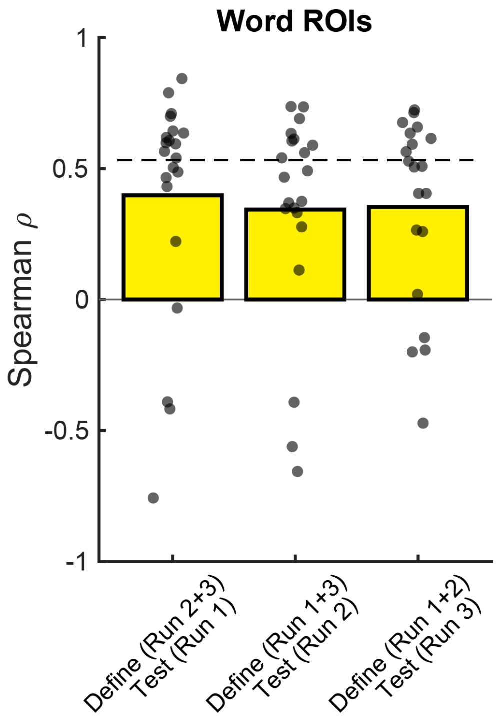


**Fig. S10 Cross-run validation analysis of OFC word patches.**
Word patches were defined on two runs ($p < 0.05$) in all 21 participants, without preselection, and tested on the remaining, independent run. Bars represent the group-averaged Spearman ρ on test runs (n = 21 for test runs 1 and 2; n = 20 for test run 3, as one participant completed only two runs). Dots represent Spearman ρ in individual participants. Dashed lines indicate uncorrected within-participant significance levels ($p < 0.05$; ρ = 0.53). In each test run, 8–12 participants showed a significant effect (estimated population prevalence = 0.37–0.55; Table S2), and the group-averaged ρ was significantly above zero (one-sample t-tests, all ps < 0.001).

**Table S3. Test of replicability of OFC word patches and prevalence summary**

| ROI Type | Define (run) | Test (run) | First-level Prevalence | MAP | HPDI96 Lower | HPDI96 Upper |
|---|---|---|---|---|---|---|
| word | 2+3 | 1 | **12/21** | 0.55 | 0.32 | 0.76 |
| word | 1+3 | 2 | **9/21** | 0.40 | 0.19 | 0.62 |
| word | 1+2 | 3 | **8/20** | 0.37 | 0.16 | 0.60 |

ROIs were defined on two runs ($p < 0.05$; n = 21) and tested on the held-out run (Spearman rho; first-level significance: rho > 0 and $p < 0.05$). One participant had only two runs: ROI defined on run 2 and tested on run 1, and ROI defined on run 1 and tested on run 2; this participant is therefore not included in the test-run-3 row (n = 20).

### OFC signal measurement and drop-out

Because the OFC is susceptible to signal dropout, we evaluated data quality in each ROI using tSNR. Both datasets were acquired with the same sequence and slab positioning, optimized to cover the OFC while minimizing dropout near the ear canals in the VOTC (see Methods).

In Dataset 1, tSNR in OFC face and color patches (38.2 and 38.6) was comparable to that of VOTC face (39.4–47.7) and color patches (36.8–43.3), in particular to the middle and anterior VOTC patches, and did not differ between runs 4 and 5 (paired t-test, $p = 0.23$; Fig. S11A). In a representative participant, tSNR was reduced in parts of the medial OFC but remained high in the lateral OFC, where the category-specific patches were located (Fig. S11B). tSNR in the OFC patches did not differ between aphantasic and typical imagers (OFC-f: 36.1 vs. 40.3, $t(18) = -1.42$, $p = 0.17$; OFC-c: 39.2 vs. 38.1, $t(18) = 0.45$, $p = 0.66$); among all ROIs, only OFA showed lower tSNR in aphantasic individuals (42.3 vs. 53.0, $t(18) = -3.54$, $p = 0.002$). In Dataset 2, tSNR in OFC word patches (35.8–35.9) was also comparable to that of fusiform word patches (39.6-40.7) and stable across the three runs (Fig. S11C).

Thus, the OFC patches retained sufficient signal quality for reliable estimation of the reported effects.

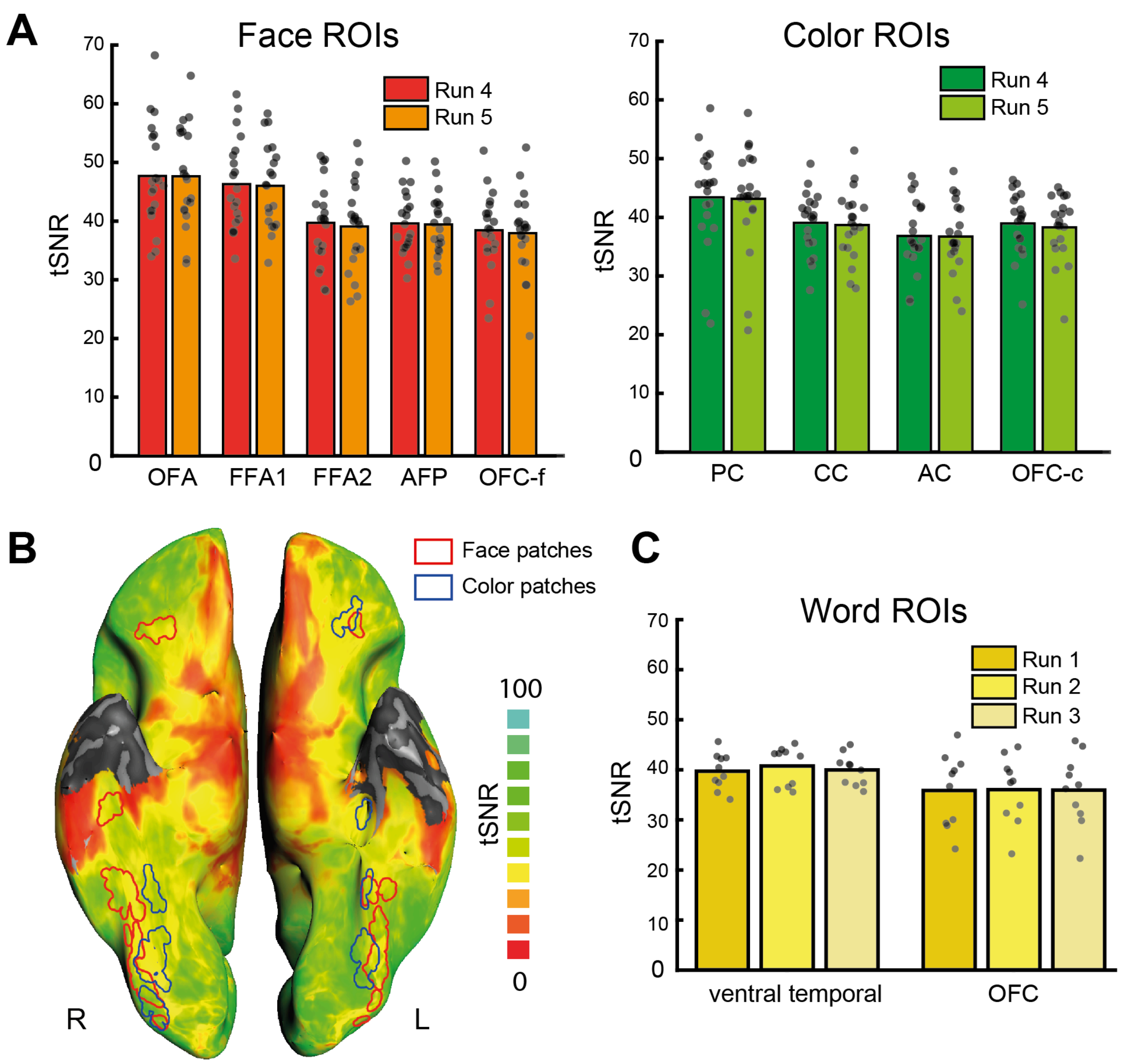


**Fig. S11 Temporal signal-to-noise ratio (tSNR) in category-specific ROIs.**
**A.** tSNR in face (left) and color (right) volume ROIs in Dataset 1 for run 4 and run 5 (439 volumes per run, TR = 2 s). Bars represent group averages (N = 20) and dots represent individual participants.
**B.** tSNR map of one representative participant, projected onto the cortical surface for visualization purposes (ventral view; unsmoothed). Red and blue contours denote face- and color-specific patches, respectively. For visualization, volumetric ROIs were dilated to compensate for sampling loss when projected onto the surface. Gray areas indicate regions outside the acquisition slab; the temporal poles were excluded to reduce susceptibility-related dropout near the ear canals.
**C.** tSNR in anterior fusiform and OFC word volume ROIs in Dataset 2 for runs 1–3 (399 volumes per run). Bars represent group averages and dots represent individual participants (n = 10).
(End)
